# Site-specific genome engineering of primary human natural killer cells for programmable anti-tumor function

**DOI:** 10.1101/2025.10.05.680386

**Authors:** Vincent Allain, Allison G. Rothrock, Pierre-Louis Bernard, William A. Nyberg, Alexis Talbot, Joseph J. Muldoon, Jing-Yi Chung, Angela To, Christopher R. Chang, Gabriella R. Kimmerly, Chang Liu, Tasha Tsao, Yasaman Mortazavi, Jin Seo, Zhongmei Li, Avishai Shemesh, Ralf Schmidt, Carl C. Ward, Alexander Marson, Lewis L. Lanier, Oscar A. Aguilar, Justin Eyquem

## Abstract

Natural killer (NK) cells are emerging as a promising platform for engineered adoptive cell therapies. However, gene editing in NK cells remains challenging, and more effective strategies are needed. Here, we established a robust, feeder-free, and modular workflow for genome engineering in primary human NK cells, combining CRISPR/Cas9 with AAV6-mediated transgene delivery. Efficient site-specific transgene integration was achieved at various loci and can be coupled with concurrent disruption of the target locus in a single editing step. Furthermore, transgene expression was tunable according to the integration site and promoter. We applied this strategy to target a chimeric antigen receptor (CAR) transgene to a panel of inhibitory NK receptor loci, establishing a synergistic approach to enhance anti-tumor activity and facilitate the reliable comparison of CAR variants without expression bias. We identified *TIGIT* as an ideal locus that supports strong CAR expression and anti-tumor function. This genome engineering framework, which leverages multiple, complementary and precisely controlled genetic edits, can support the rational design of future NK-cell therapies tailored to overcome cell-intrinsic limitations and tumor-specific barriers.

## Main text

Chimeric antigen receptor (CAR) T cells have revolutionized the clinical management of patients with hematological malignancies^1^. Advances in gene editing are driving the development of next-generation CAR T cells to overcome biological hurdles and broaden their clinical indications^2–5^. More recently, natural killer (NK) cells, which exhibit potent innate anti-tumor activity^6–9^, have emerged as an alternative cell type for CAR-mediated tumor targeting^10,11^. NK cells offer several potential advantages as compared to T cells, particularly in the allogeneic setting as they do not cause graft-versus-host disease^12–17^. However, despite promising early clinical successes and a favorable safety profile^18–22^, CAR NK cells remain a less advanced approach, and their development is hindered by both technical and biological challenges. We hypothesized that CAR NK cells, like their T-cell counterparts, would benefit from gene editing techniques, namely the targeted integration of a CAR transgene at defined genomic loci rather than the semi-random integration obtained using gammaretroviral (gRV) vectors. Precise targeting of a CAR to the *TRAC* locus has proven advantageous for the efficacy of CAR T cells^23^. Furthermore, disruption of NK inhibitory receptors via functional blockade or genomic deletion appears to be a promising path for enhancing anti-tumoral activity^24–26^.

Here, we report a feeder-free, optimized gene editing workflow for primary human NK cells allowing the evaluation of the targeted integration of a CAR transgene at key NK cell-relevant loci. We also demonstrate that the targeted integration of a CAR transgene can be combined with simultaneous ablation of inhibitory receptors in one editing step. Finally, we focus on the inhibitory receptor TIGIT as a model integration locus to compare different exogenous promoters and CAR variants. Altogether, we present a versatile workflow for next-generation CAR NK cell engineering.

## Results

### Optimization of a feeder-free gene ablation protocol for efficient editing of primary NK cells

We followed a strategy of stepwise improvement starting from a previously published feeder-free protocol^27^ (**Fig. 1a**). We optimized CRISPR-Cas9-mediated gene disruption using *CD38* as a model locus, as CD38 ablation in NK cells has been reported to prevent their depletion in the context of anti-CD38 monoclonal antibody treatment^28,29^. Two sgRNAs targeting *CD38* exon 1 were each tested with the electroporation pulse code DN-100 in a Lonza 4D nucleofector (**Fig. 1b,c**). The higher-efficiency sgRNA (gRNA2) was selected for further use. Similar knockout (KO) efficiency was achieved despite increasing the number of electroporated cells from 0.5 x 10^6^ to 2 x 10^6^ per well for a fixed quantity of RNP (40 pmol) (**Supplementary Fig. 1**). In comparing the baseline electroporation pulse code DN-100^27^ to the previously reported CM-137^30^, we observed similar KO efficiency but higher viable cell recovery (47% median increase) at day 4 post-electroporation with CM-137, which we used for further experiments (**Fig. 1d,e,f,g**). Notably, high KO efficiency was replicated at the *B2M* locus (**Fig. 1h,i,j**). Our workflow was readily adaptable for *CD38* ablation in NK cells transduced with a RD114-pseudotyped gRV vector^31^ for CAR expression (**Fig. 1k,l**). Finally, we demonstrated the versatility of our protocol through efficient base-editing using a mRNA-delivered adenine base editor (ABE) targeting a splice site in the *CD7* locus and resulting in *CD7* KO^32^, as validated by flow cytometry and next-generation sequencing (**Fig. 1m,n,o,p**). Notably, CD7 disruption has been reported as a clinically relevant strategy to prevent fratricide in the context of anti-CD7 CAR therapies^33,34^.

**Figure 1.**
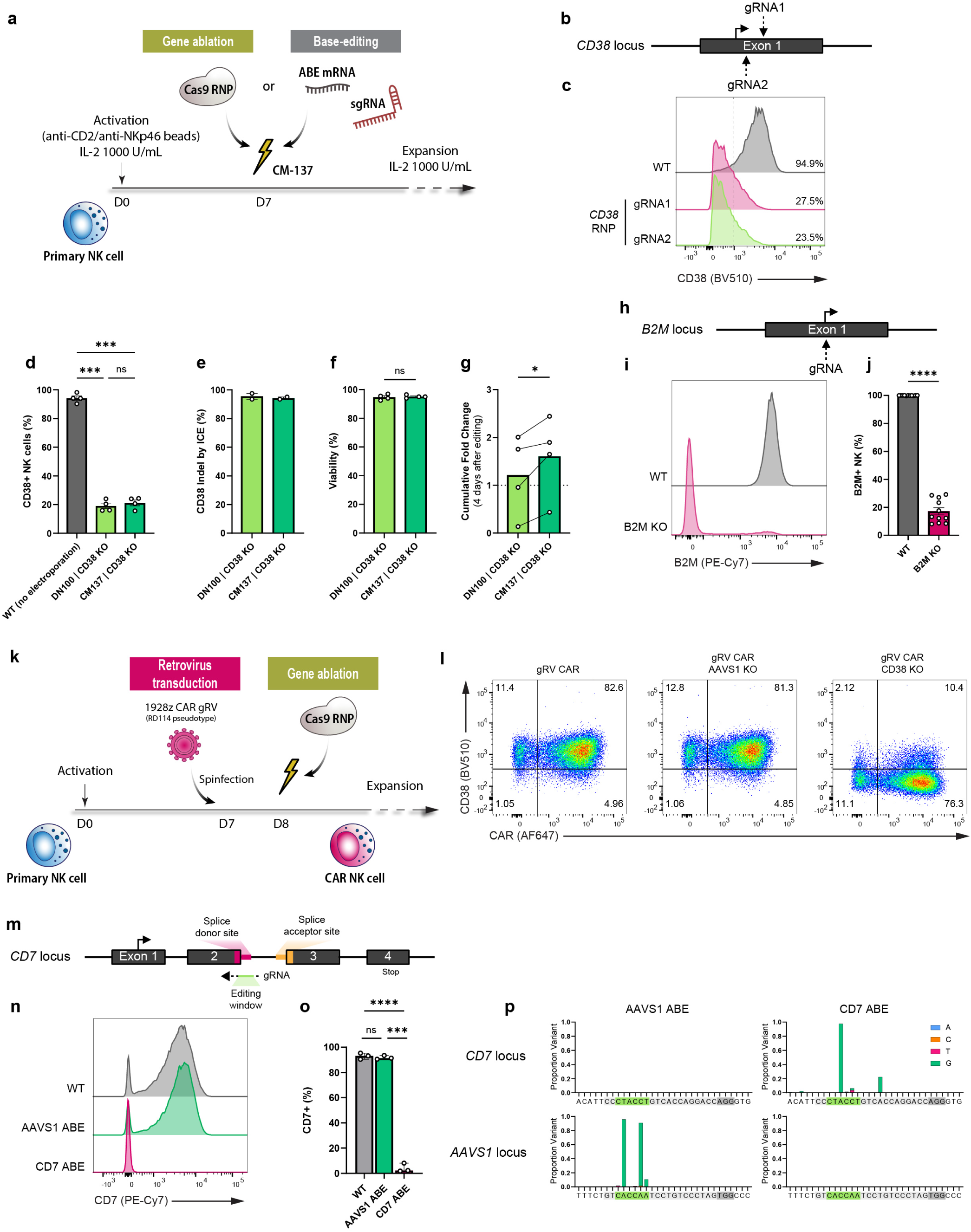
Feeder-free editing strategy for efficient gene ablation and base-editing in primary NK cells. **a**, Schematic of feeder-free editing strategy for gene ablation or base-editing in primary NK cells. **b**, Representation of the cut sites at *CD38* locus for the screening of 2 sgRNAs. **c**, Representative flow cytometry histogram for CD38 expression assessed 4 d after editing (pulse code DN-100). Guide gRNA3 was used for subsequent panels. **d-g**, Comparison of outcomes assessed 4 d after editing for NK cells electroporated using two electroporation pulse codes (DN-100 versus CM-137): editing efficiency assessed by flow cytometry (**d**), indel percentage assessed at the genomic level by ICE analysis (**e**), viability assessed by trypan blue staining on a Countess II counter (**f**) and total viable cell recovery from editing (**g**). *n*=4 biological replicates from different human donors except for (**e**) where *n*=2. **h-j**, Validation of the gene ablation workflow by targeting the exon 1 of *B2M* (**h**), assessed by flow cytometry (**i,j**). *n*=11 biological replicates from distinct human donors. **k**, Schematic of feeder-free gammaretroviral (gRV) transduction combined with CRISPR-mediated gene ablation. **l**, Representative flow cytometry data showing the expression of CD38 and CAR for gRV transduced CAR NK, without or with subsequent gene ablation (*AAVS1* as a control or *CD38*). **m-p**, Extension of the feeder-free editing workflow to base-editing by co-electroporating an adenine base editor (ABE) mRNA with a sgRNA. A splice donor site of *CD7* was targeted for base-editing (**m**), resulting in a loss of CD7 expression assessed by flow cytometry compared to WT NK cells or NK cells edited at the *AAVS1* locus (**n,o**). *n*=3 biological replicates from different human donors. Efficiency of specific adenine base-editing was confirmed at the genomic level for *AAVS1* and *CD7* base editing in 2 independent donors (x-axis: genomic WT sequence, light grey: gRNA, dark grey: PAM, green: theoretical editing window) (**p**). Data are shown as mean ± SEM. *P*-values are from repeated-measures one-way ANOVA with Tukey’s multiple comparison tests (**d,o**) and from a two-tailed paired *t* test (**f,g,j**). *, *p* ≤ 0.05; **, *p* ≤ 0.01; ***, *p* ≤ 0.001; ****, *p* ≤ 0.0001. ns, not significant.

### Efficient gene targeted integration using AAV6 to deliver an HDRT

To provide a site-specific alternative to the semi-random transgene integration associated with gRV vectors, we adapted our protocol for knock-in (KI). Following RNP electroporation, a recombinant adeno-associated virus 6 (AAV6) homology-directed repair template (HDRT) was delivered for transgene integration (**Fig. 2a**). To define optimal parameters for this protocol, we used a strategy based on the in-frame integration of a promoter-less green fluorescent protein (GFP) sequence at the start of the coding sequence of the clathrin A (*CLTA*) gene. Upon targeted integration, the resulting GFP-CLTA fusion gene is expressed under the control of the endogenous *CLTA* promoter^35,36^ (**Fig. 2b**). As expected, no episomal expression was observed when NK cells were transduced with AAV alone, consistent with reliance on the endogenous promoter for expression (**Fig. 2c**). In contrast, strong GFP expression (up to 60% of cells) was detected by flow cytometry when *CLTA* RNP electroporation was paired with AAV transduction, reflecting GFP KI. We compared the electroporation pulse codes CM-137 and DN-100 (**Fig. 2d,e,f,g**) and observed, despite similar KI efficiency and viability 4 days after editing, that the overall yield of edited cells was 4.8-fold higher (median) with CM-137. When removing human platelet lysate (HPL) from the culture media during AAV transduction, we observed a negative impact of HPL on KI efficiency, an effect already known with serum and neutralizing antibodies on AAV6 transduction^37,38^ (**Fig. 2c** and **Extended Data Fig. 1a,b,c**). This effect was evident primarily at low multiplicity of infection (MOI) and was overcome by using higher MOI. However, the potential benefit of removing HPL during transduction was counterbalanced by increased toxicity, as indicated by absolute cell counts. Importantly, NK cells were able to recover and proliferate after editing (**Extended Data Fig. 1d**). In summary, high KI efficiency was achieved using AAV6 for HDRT delivery. Among the conditions tested, combining CM-137 pulse code with HPL maintenance during AAV transduction provided the best balance between KI efficiency and cell viability, resulting in an optimal yield.

**Figure 2.**
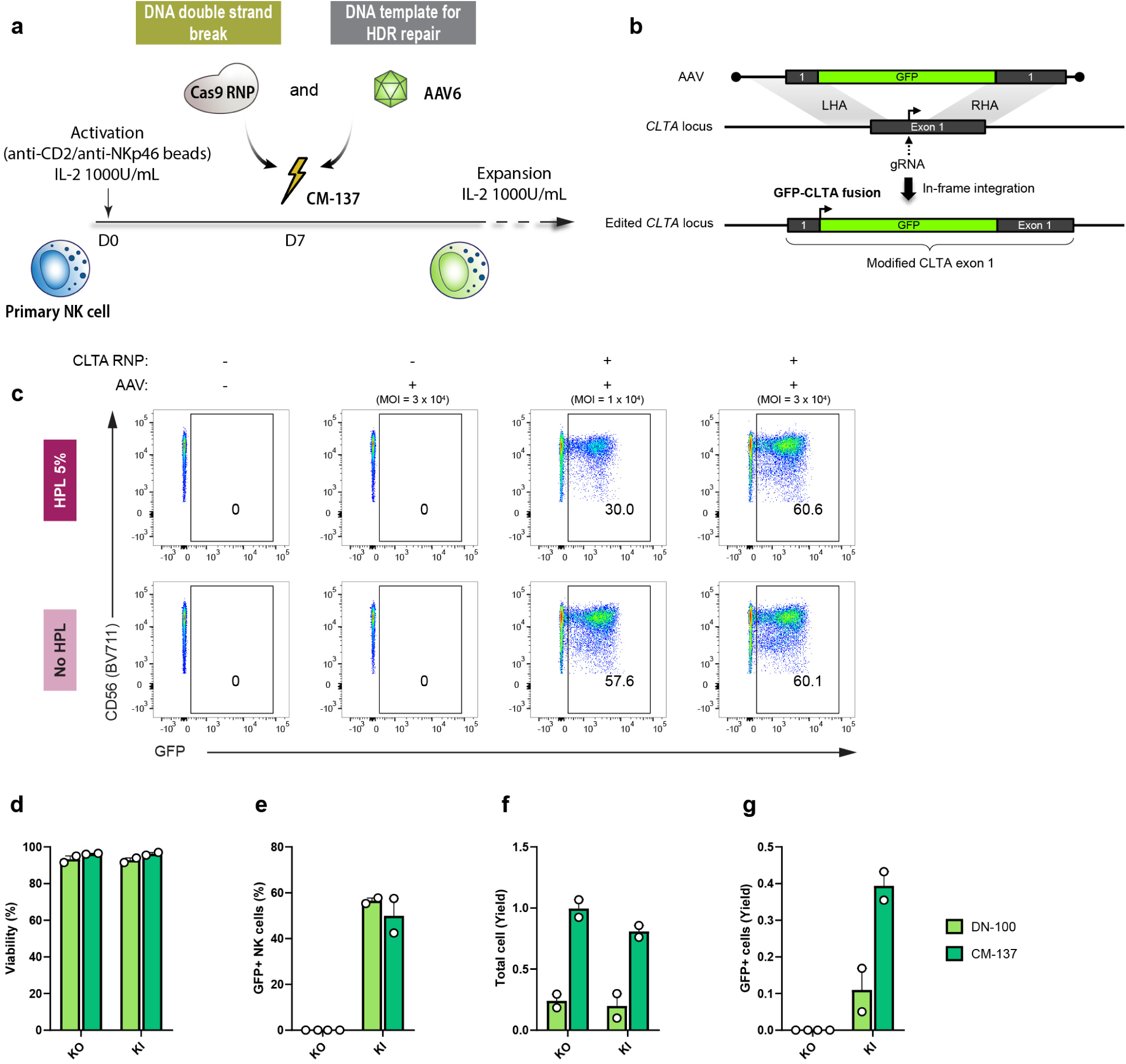
Efficient feeder-free targeted integration of a GFP fusion transgene at the *CLTA* locus in primary NK cells using CRISPR/Cas9 and AAV6. **a**, Schematic of feeder-free editing strategy for targeted integration of a transgene in primary NK cells using AAV6 to deliver an HDRT. **b**, Schematic depiction of the gene editing strategy for GFP targeted integration at the *CLTA* locus. The HDRT allows in-frame integration of a promoter-less GFP fusion tag at the start of the coding sequence of *CLTA*, resulting in a GFP-CLTA fusion protein replacing the native CLTA protein. **c**, Representative flow cytometry data showing GFP expression after GFP targeted integration in primary NK cells and the impact of HPL in the culture media during AAV6 transduction. **d-g**, Pulse codes DN-100 and CM-137 were compared for editing outcomes assessed 4 d post-editing. Viability was evaluated by trypan blue staining on a Countess II counter (**d**), and the percentage of GFP^+^ NK cells (**e**) was assessed by flow cytometry. Cumulative fold-change since editing for the total cell number (**f**) and the number of GFP^+^ NK cells (**g**) are represented. *n*=2 biological replicates from different human donors in 2 different experiments.

### Generation of CAR NK cells by targeting a CAR to the *B2M* locus

The optimized editing protocol was applied to generate a more clinically relevant NK cell product through site-specific CAR transgene integration. An anti-CD19 CAR with the CD28 costimulation domain (1928z) was integrated at *B2M* exon 1 (sgRNA targeting the start codon) under the regulation of the *B2M* endogenous promoter (**Fig. 3a**). We previously reported a similar strategy in primary T cells, resulting in high and homogeneous CAR expression^23^. Given the innate recognition and targeting by NK cells of cells with low or absent HLA class I expression (“missing self”)^39,40^, we hypothesized that an editing strategy preserving B2M expression upon CAR integration would be advantageous by preventing fratricide of *B2M*-CAR NK cells and also driving spontaneous enrichment of the B2M-rescued CAR^+^ cell fraction, as *B2M* KO cells without CAR integration would become depleted (**Fig. 3b**). Therefore, we designed two HDRTs differing by the disruption or rescue of the B2M coding sequence upon targeted integration (**Fig. 3a**). As shown previously (**Fig. 1h,i,j**), *B2M* KO was highly efficient, and transduction with a recombinant AAV6 to deliver the HDRT resulted in efficient generation of *B2M*-CAR NK cells (up to 80%) (**Fig. 3c** and **Extended Data Fig. 2f**). Using the *B2M*-CAR KI rescue design, we demonstrated that the use of an NHEJ inhibitor (M3814) at a concentration of 0.5 µM further enhanced CAR KI efficiency (**Supplementary Fig. 2a,b,c**). CAR expression was higher with the *B2M* rescue design compared to the *B2M* disruptive design, suggesting that the endogenous 3’ UTR drove a higher transcriptional level (**Fig. 3c** and **Extended Data Fig. 2a,b,c,d**). As expected, B2M expression was ablated in the disruptive design, and the rescue design led to a partial expression of B2M upon CAR integration (**Fig. 3c**). Despite intermediate B2M expression, the *B2M* rescue design was effective at preventing fratricide and promoted a progressive enrichment of the CAR+ fraction (**Fig. 3d** and **Extended Data Fig. 2f**). By contrast, we observed a progressive loss of B2M^-^ cells (both CAR^-^ and CAR^+^) in the *B2M* KO and the *B2M*-CAR KI (*B2M* disruptive design) conditions (**Fig. 3d** and **Extended Data Fig. 2e**).

**Figure 3.**
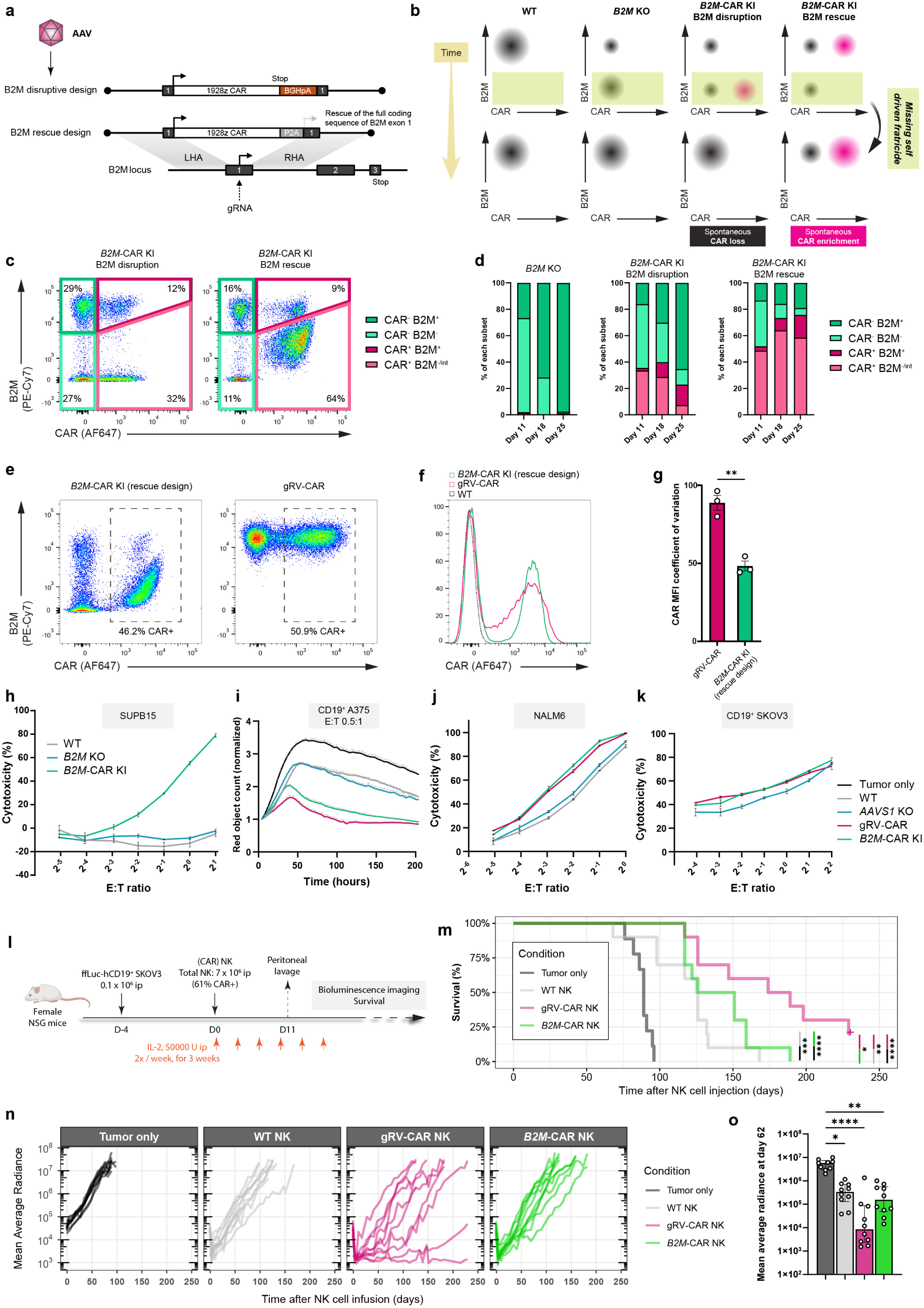
Targeting a CAR to the *B2M* locus in primary NK cells for enhanced anti-tumor activity. **a**, Schematic depiction of the editing strategy for targeted integration of a 1928z CAR at the *B2M* locus (*B2M* disruptive and *B2M*-rescue HDR templates). **b**, Concept of spontaneous CAR+ NK cell enrichment by leveraging missing-self driven fratricide. Upper panels represent the theoretical starting point after editing for the different conditions, lower panels represent the expected evolution over time **c**, Representative flow cytometry data of primary NK cells edited using the transgenes shown in (**a**) assessed 4 d after editing, and definition of the subpopulations used in panel (**d**). **d**, Representative evolution over time of the different cell subsets based on CAR and B2M expression for *B2M* KO NK and *B2M*-CAR NK cells (B2M-disruptive and B2M-rescue designs). **e-g**, Comparison of *B2M*-CAR KI NK (rescue design) and NK cells transduced by a gRV vector for CAR integration (gRV-CAR). Representative flow cytometry dot plots (**e**) and histograms (**f**) showing the respective B2M and CAR expressions across conditions, and corresponding CAR MFI coefficient of variation. *n*=3 donors (3 independent experiments). Data are shown as mean ± SEM. *P*-values are from a two-tailed paired *t*-test. **h**, Luciferase-based cytotoxicity assay against the naturally resistant SUPB15 cell line (E:T ratio based on the number of CAR^+^ NK cells). **i**, Incucyte cytotoxicity assay against CD19^+^ A375 cell line, at 0.5:1 E:T ratio (based on the total number of NK cells). **j-k**, Luciferase-based cytotoxicity assay against NALM6 (**j**) and CD19^+^ SKOV3 (**k**) cell lines (E:T ratio is based on the total number of NK cells). **l**, Schematic experimental design of the SKOV3 xenograft mouse model for *in vivo* assessment of CAR NK anti-tumor efficacy. **m**, Kaplan-Meier curves for survival analysis. *P*-values are from a log-rank test. **n-o**, Mean average radiance assessed by bioluminescence imaging across the course of the experiment (**n**) and at d 62 (**o**). Data are shown as median ± IQR. *P*-values are from Kruskal-Wallis test with Dunn’s multiple comparison tests against the tumor only condition. *, *p* ≤ 0.05; **, *p* ≤ 0.01; ***, *p* ≤ 0.001; ****, *p* ≤ 0.0001.

The CAR and B2M expression pattern was similar when the strategy was applied to T cells (**Supplementary Fig. 3a**). To investigate whether the diminished B2M expression observed with the rescue design was associated with monoallelic KI, we co-transduced two AAVs (one HDRT for a CAR and the other for a truncated EGFR (EGFRt), both with a *B2M* rescue strategy) (**Supplementary Fig. 3b**). B2M expression was diminished even in the population of cells co-expressing the CAR and EGFRt, demonstrating that *B2M* haplo-insufficiency did not explain the lower B2M expression level (**Supplementary Fig. 3c,d,e**). This result also demonstrated that our editing protocol led to efficient biallelic KI (estimated as >50%).

### *In vitro* anti-tumor activity of *B2M*-CAR NK is not recapitulated *in vivo* in a solid tumor model

As compared to NK cells transduced with a gRV vector encoding the same CAR construct (gRV-CAR NK cells), *B2M*-CAR NK cells (rescue design) exhibited significantly more homogeneous CAR expression (**Fig. 3e,f,g**). Functionally, *B2M*-CAR NK cells exhibited *in vitro* cytotoxic activity against the SUPB15 B-cell leukemia cell line which is naturally CD19^+^ and normally resistant to NK cell killing (**Fig. 3h**). *B2M*-CAR NK cells also were cytotoxic against different CD19^+^ target cell lines (NALM6 B-cell leukemia cell line, CD19-transduced A375 melanoma cell line, CD19-transduced SKOV3 ovarian adenocarcinoma cell line), with similar cytotoxicity compared to gRV-CAR NK cells (**Fig. 3i,j,k**). *B2M*-CAR NK and gRV-CAR NK cells were compared using a solid tumor xenograft model (**Fig. 3l**). Briefly, female NSG mice engrafted intraperitoneally with CD19^+^ SKOV3 human ovarian cancer cells were treated with a single intraperitoneal (i.p.) injection of NK cells, followed by twice-weekly i.p. injections of interleukin-2 (IL-2). Assessment of tumor burden by bioluminescence imaging (BLI) revealed a rapid anti-tumor response for all the treatment conditions (WT NK, gRV-CAR NK and *B2M*-CAR NK) (**Fig. 3n,o**) with deeper initial control by CAR-expressing NK cells (**Extended Data Fig. 3a,b**), resulting in an increased overall survival as compared to untreated mice (**Fig. 3m**). However, most of the mice eventually relapsed. Only the gRV-CAR NK cell condition had significantly delayed relapses, which conferred a significant survival advantage compared to both WT NK and *B2M*-CAR NK conditions (**Fig. 3m**). A predetermined subset of mice (n=4 per condition) was euthanized on day 11 and a peritoneal lavage was performed (**Extended Data Fig. 3a,c**). NK cell recovery was highly variable across conditions, with no group demonstrating a clear advantage in persistence or proliferation. Notably, the proportion of CAR^+^ cells in the gRV-CAR NK group was similar to the infused product, whereas the *B2M*-CAR NK group had a slight but significant increase (**Extended Data Fig. 3d,e,f,g,h**). While providing high and reproducible transgene expression, these results suggest that the *B2M* locus may be suboptimal for NK cell engineering and underscore the need to explore alternative integration loci.

### Screening NK-relevant loci for targeted integration of a CAR transgene identifies *TIGIT* as a promising locus

Building on previous results, we expanded our editing approach to screen several NK-relevant loci for the integration of a CAR transgene. The primary goal was to identify a set of loci permitting high KI efficiency and high CAR expression. For loci encoding *NCAM* (CD56) and NK activating receptors (*FCGR3A*, CD16A; *NCR1,* NKp46; *NCR3*, NKp30; and *CD226*, DNAM-1), we preserved the receptor expression and used the endogenous promoter. Therefore, a non-disruptive KI design similar to the one described at *B2M* was favored. Conversely, in the case of loci encoding NK inhibitory receptors (*KLRC1*, NKG2A; *KLRB1*, CD161; *TIGIT*; and *LILRB1*), we used the CAR KI to concurrently disrupt the targeted receptor, a KO by KI strategy that we expected could provide a synergistic anti-tumor effect. We opted for a reverse integration strategy for the CAR transgene with respect to the target gene, in combination with the use of the elongation-factor 1 α subunit (EF1α) exogenous promoter^41^ (**Fig. 4a,b** and **Extended Data Fig. 4a,b**). For each screened locus, we designed and validated sgRNAs with high KO efficiency – except for *LILRB1* where the efficiency remained limited (**Supplementary Fig. 4a,b**). After delivering the corresponding HDRT, we observed variable frequencies of CAR^+^ cells, reaching up to 70% for *KLRC1*, 67% for *KLRB1,* and 47% for *NCAM* at four days after editing. CAR expression levels varied across loci (**Fig. 4c,d,e**), with the highest levels obtained at *KLRC1*, *NCAM*, *KLRB1*, and *TIGIT*. Of note, all targeted loci yielded lower CAR expression levels compared to the *B2M* locus. For activating receptors, the rescued receptor was only partially expressed, as observed with *B2M* (**Extended Data Fig. 5**). We selected the top-ranked CAR NK conditions in terms of CAR^+^ percentage (*KLRC1*, *KLRB1*, *NCAM*, *TIGIT*, *FCGR3A*, and *NCR1*) and compared their cytotoxic function against the NALM6 and SUPB15 cell lines. Notably, all the conditions performed better than the *B2M*-CAR NK cells, with *KLRC1*-CAR NK performing the best (**Fig. 4f,g**). Because the anti-tumor activity of CAR NK cells will be the net balance of multiple receptor/ligand interactions, including the impact of the potential KO of the integration locus in addition to the CAR expression level, we characterized the main NK ligands on several target cell lines to facilitate the interpretation of functional results (**Extended Data Fig. 6a,b,c**).

**Figure 4.**
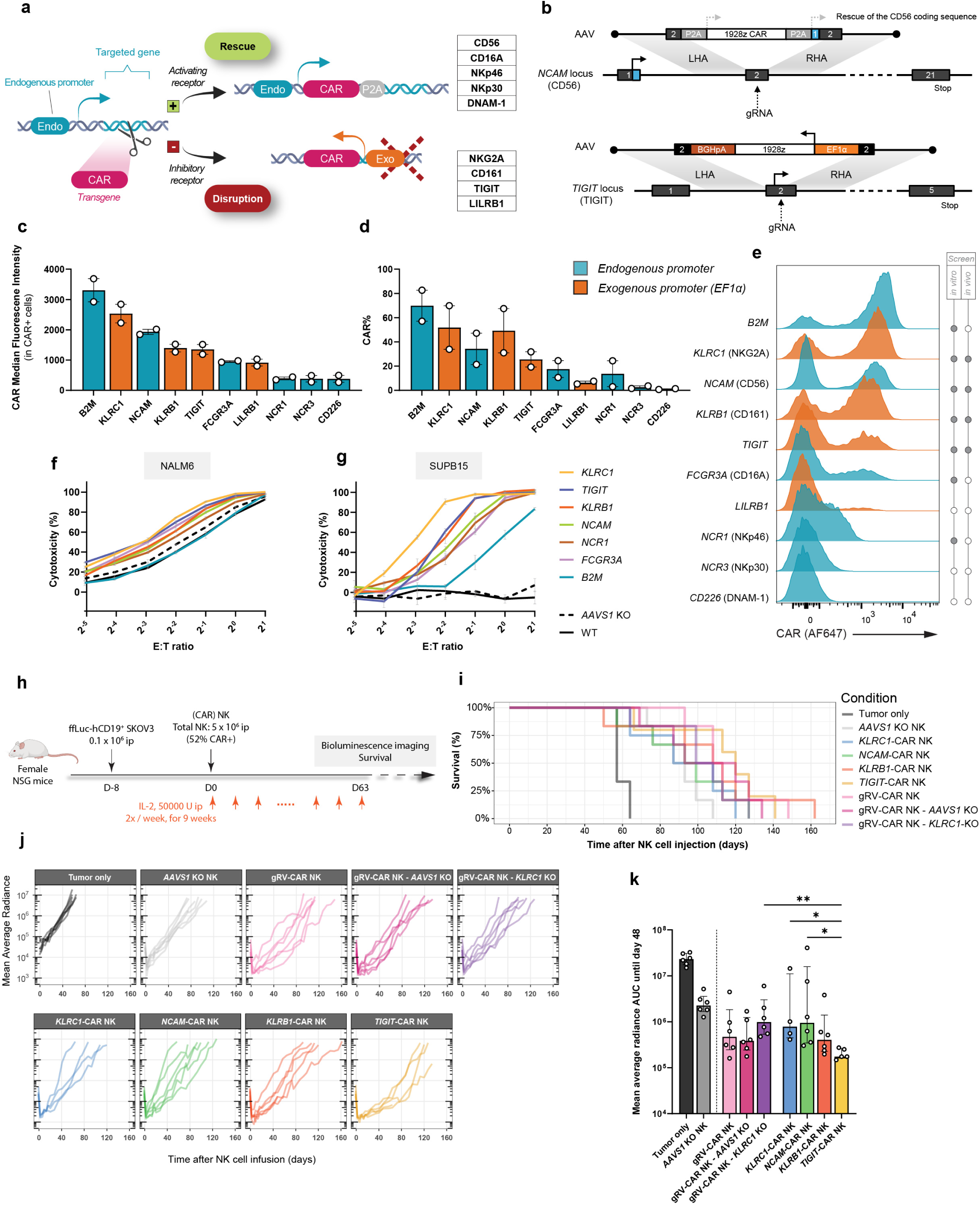
Targeting a CAR into multiple NK-relevant loci in primary NK cells identifies *TIGIT* as a promising integration locus for enhanced anti-tumor activity. **a**, Schematic of the general editing strategy for various NK-relevant loci. A rescue design leveraging the endogenous promoter of the targeted locus is used for activating receptors, whereas a disruptive design is used for inhibitory receptors, under the transcriptional control of the exogenous EF1α promoter. **b**, Schematic depiction of the gene-editing strategy and the transgene design for targeted integration of a 1928z CAR at the *NCAM* locus (activating receptor, rescue design) and at the *TIGIT* locus (inhibitory receptor, disruptive design). **c-e**, Flow cytometry evaluating CAR expression was performed 11 d after editing for 2 donors (2 independent experiments). CAR Median Fluorescence Intensity for the CAR^+^ cells is shown (**c**) as well as the corresponding KI efficiency evaluated as the percentage of CAR^+^ cells (**d**). Representative histograms of CAR expression are shown for the different editing designs (**e**). Endogenous promoter/rescue designs (blue) were distinguished from exogenous/disruptive designs (orange). **f**, Luciferase-based cytotoxicity assay against the NALM6 cell line, with E:T ratio based on the total number of NK cells. **g**, Luciferase-based cytotoxicity assay against the SUPB15 cell line, with E:T ratio based on the number of CAR^+^ NK cells. **h**, Schematic experimental design of the SKOV3 xenograft mouse model for *in vivo* assessment of CAR NK anti-tumor efficacy. **i**, Kaplan-Meier curves for survival analysis. *P*-values from a log-rank test are described in **Supplementary Fig. 5. j**, Mean average radiance assessed by bioluminescence imaging across the course of the experiment. **k**, Area under the curve of the BLI mean average radiance until d 48 (last measurement day on which all mice were alive). Data are shown as median ± IQR. *P*-values are from Kruskal-Wallis test with Dunn’s multiple comparisons test comparing each CAR NK condition to the *TIGIT*-CAR NK condition used as a reference.

The four loci with the highest KI efficiency and strong *in vitro* cytotoxicity (*KLRC1*, *KLRB1*, *NCAM*, and *TIGIT*) were evaluated *in vivo* using the CD19^+^ SKOV3 xenograft model (**Fig. 4h**). NK cells with a KO at the safe harbor locus *AAVS1* were used as a control to account for the effect of electroporation and DNA double-strand break. In addition, we included retrovirally transduced CAR NK cells (gRV-CAR NK), some of which were ablated for *KLRC1* (our top candidate locus based on *in vitro* data) or *AAVS1*. We observed a rapid and profound anti-tumor response after the injection of NK cells for all conditions. Notably, *TIGIT*-CAR NK cells significantly enhanced tumor control during the initial period up to day 48, the last time point with BLI data when all mice were still alive (**Fig. 4j,k**). *TIGIT*-CAR NK cells also conferred significantly increased overall survival compared to *AAVS1* KO NK cells (**Fig. 4i** and **Supplementary Fig. 5a,b,c**). Interestingly, while *KLRC1* KO was best in the *in vitro* cytotoxicity assay, both *KLRC1*-CAR NK (KI) and *KLRC1* KO gRV-CAR NK cells performed poorly compared to *AAVS1* KO gRV-CAR NK cells in this *in vivo* model. In summary, *in vivo* evaluation using a xenograft model of ovarian cancer identified *TIGIT* as a favorable genomic integration site for generating potent CAR NK cells.

### Tuning CAR expression at the *TIGIT* locus

Transgene KI allows for the unbiased comparison of the strength of different exogenous promoters. We compared five exogenous promoters to control the expression of a CAR transgene integrated at the *TIGIT* locus. We screened the ubiquitin C (UbC) promoter, phosphoglycerate kinase (PGK) promoter, enhancer sequence from the SFG gammaretrovirus used in our work (Mo-MLV LTR, referred to as LTR here)^42^, and MND promoter^43,44^, in addition to the EF1α promoter used previously (**Extended Data Fig. 7a**). The promoter series provided a wide range of CAR expression. The highest level was observed with EF1α, followed by PGK (a 4-fold reduction in expression). Ubc and retroviral-derived promoters (MND and LTR/SFG) yielded the lowest expression levels, >10-fold less than with EF1α (**Extended Data Fig. 7b,c,d**). Overall, we demonstrated the feasibility of fine-tuning CAR expression in primary NK cells by using different exogenous promoters and identified EF1α as the most potent among those evaluated.

### Calibration of CAR signaling using targeted integration of ITAM variants at the *TIGIT* locus

Given the importance of ITAM-bearing chains in NK receptor signaling and the demonstration that CAR signaling can be fine-tuned in CAR T cells by using ITAM variants of the CD3ζ domain^45^, we compared three ITAM variants of the 1928z CAR in primary NK cells: the regular 123 CAR (the 3 ITAMs are functional), the 1XX variant (ITAMs 2 and 3 are inactivated by Y-to-F substitutions) and the XX3 variant (ITAMs 1 and 2 are inactivated similarly) (**Extended Data Fig. 8a**). *TIGIT* KI was used to reduce biases related to differences in CAR expression (**Extended Data Fig. 8b,c**). Of note, we incorporated a compact CD34 epitope tag adapted from the Q8 marker^46^ in a bicistronic fashion with the CAR. We did not observe any differences in the anti-tumor *in vitro* activity against NALM6, CD19^+^ SKOV3, and CD19^+^ A375 (**Extended Data Fig. 8d,e,f**). However, enhanced anti-tumor activity was observed with the 1XX variant against the SUPB15 cell line (**Extended Data Fig. 8g**). We combined the targeted integration of the CAR variants at *TIGIT* with the retrovirus-mediated delivery of a transgene encoding soluble IL-15 (sIL-15) and an EGFRt marker (**Fig. 5a** and **Extended Data Fig. 9a**), which we validated functionally *in vitro* (**Extended Data Fig. 9b-h**). Constitutive expression of sIL-15 is a strategy used previously, including in clinical trials, that aims to promote NK cell proliferation and persistence^20,47,48^. The combined editing strategy yielded NK cells with similar expression levels of both CAR and sIL-15/EGFRt under all conditions (**Fig. 5b**). The same *in vitro* cytotoxicity results were replicated with the constitutive expression of sIL-15, and degranulation and cytokine production were in alignment with the cytotoxicity data (**Supplementary Fig. 6a-i**). Furthermore, we did not observe phenotypic differences between the ITAM variants at steady state (**Supplementary Fig. 7a**). We tested these ITAM variants in two *in vivo* models. First, the aggressive acute lymphoblastic leukemia NALM6 model, widely used in the context of CAR-T therapies, demonstrated lower anti-tumor activity with the XX3 variant compared to the 123 CAR and the 1XX variant (**Fig. 5c,e,f,h**). The three ITAM variants were tested in the SKOV3 model (**Fig. 5i**). Again, the XX3 variant was detrimental compared to the 123 and 1XX variants. Peritoneal lavage was performed on all surviving animals after week 20, and NK cells were quantified and characterized phenotypically (**Supplementary Fig. 8a-l**). Remarkably, we detected engineered NK cells more than a 100 d post-injection, and the NK/SKOV3 ratio in the lavage was significantly higher for the 123 CAR compared to the 1XX variant (**Fig. 5n**). In summary, with *TIGIT*-CAR KI we identified the XX3 variant as detrimental in two different *in vivo* models despite good *in vitro* performance, and we suggest that the benefit of the 1XX variant may depend on the tumor model.

**Figure 5.**
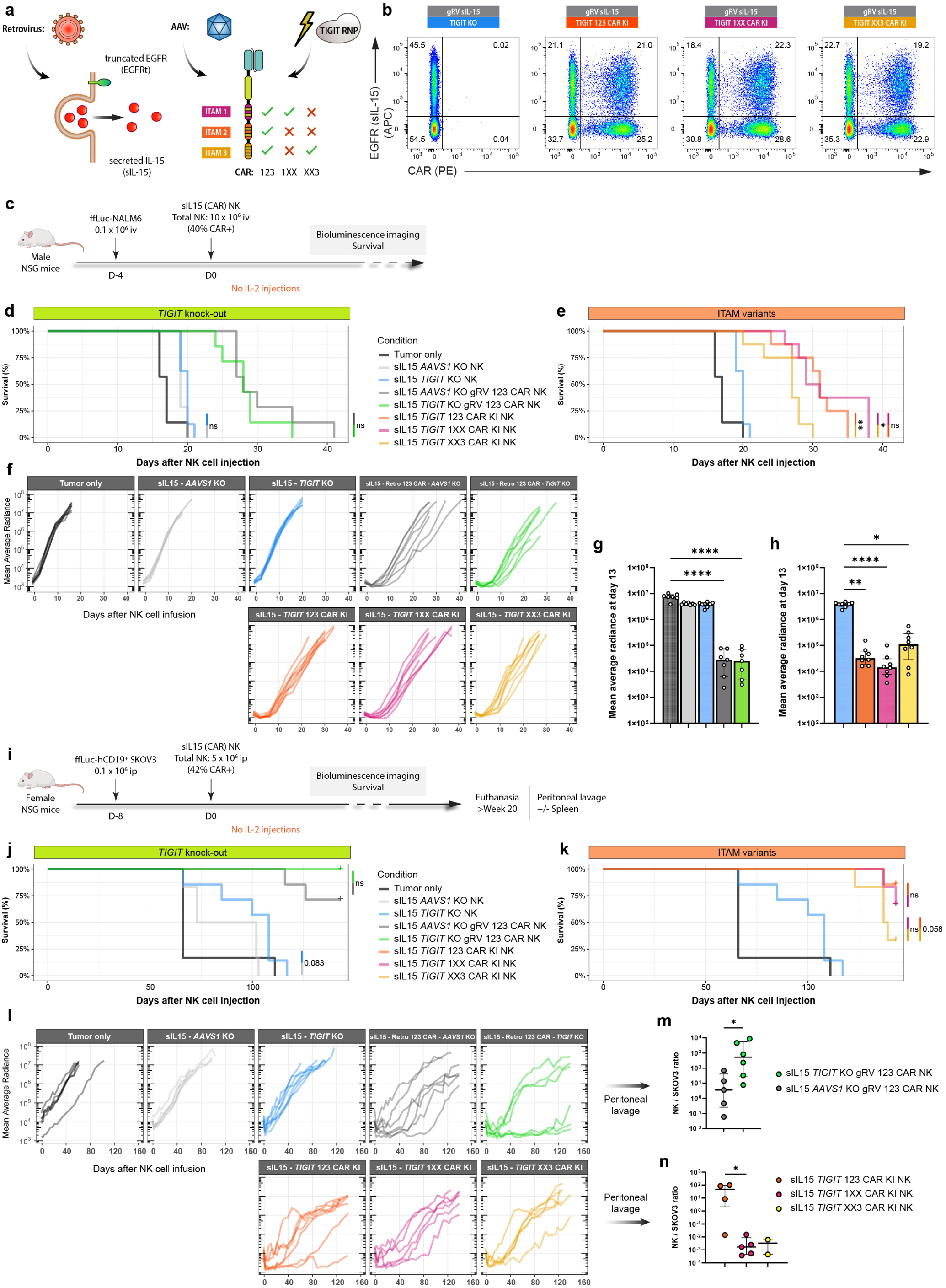
*In vivo* anti-tumor activity of CAR NK cells with ITAM variants. **a**, Primary NK cells were sequentially engineered to constitutively secrete sIL-15 (along with a EGFRt membrane marker) with a gRV vector (as detailed in **Extended Data Fig. 9a**) and to express a CAR with ITAM variants through targeted integration at the *TIGIT* locus (as detailed in **Extended Data Fig. 8a**). **b**, Representative flow cytometry data showing the editing efficiency achieved with this strategy (EGFR and CAR staining). **c**, Schematic experimental design of the NALM6 xenograft mouse model for *in vivo* assessment of CAR NK anti-tumor efficacy. **d-e**, Kaplan-Meier curves for survival analysis, with NK conditions split in 2 main groups: conditions assessing the impact of *TIGIT* KO (**d**) and conditions exploring the impact of CAR ITAM variants (**e**). P-values are from a log-rank test. **f-h**, Mean average radiance assessed by bioluminescence imaging across the course of the experiment (**f**) and at d 13 (**g-h**). Data are shown as median ± IQR. *P*-values are from Kruskal-Wallis test with Dunn’s multiple comparison tests against either the tumor only condition (**g**) or the *TIGIT* KO condition (**h**). **i**, Schematic experimental design of the SKOV3 xenograft mouse model for *in vivo* assessment of CAR NK anti-tumor efficacy. **j-k**, Kaplan-Meier curves for survival analysis, with NK conditions split in 2 main groups. *P*-values are from a log-rank test. **l**, Mean average radiance assessed by bioluminescence imaging across the course of the experiment. **m-n**, Ratio between the number of NK cells and the number of SKOV3 cells recovered from the peritoneal lavage of surviving animals after week 20, for the gRV conditions (**m**) and the ITAM variant conditions (**n**). Data are shown as median ± IQR. *P*-values are from two-tailed Mann-Whitney test. *, *p* ≤ 0.05; **, *p* ≤ 0.01; ***, *p* ≤ 0.001; ****, *p* ≤ 0.0001.

### Effect of *TIGIT* ablation on anti-tumor activity

Finally, we explored the impact of *TIGIT* ablation on the anti-tumor activity of CAR NK cells. We performed a KO of *TIGIT* or *AAVS1* in NK and gRV-CAR NK cells and did not observe phenotypic differences between conditions at steady state from *TIGIT* deletion (**Supplementary Fig. 7b**). These conditions were then tested *in vivo*. No benefit was observed with the KO of *TIGIT* in the NALM6 xenograft model (**Fig. 5c,d,f,g**), whereas a beneficial trend was observed in the SKOV3 model in terms of tumor control (including long-term controllers) and survival, both for CAR-transduced and untransduced NK cells (**Fig. 5i,j,l**). In this tumor model, NK cells and tumor cells were recovered from peritoneal lavage in surviving animals after 20 weeks (**Supplementary Fig. 8a-l**). Phenotypic characterization confirmed that the control CAR NK cells (*AAVS1* KO) continued to express TIGIT at this late timepoint (**Supplementary Fig. 8i**). Notably, a significantly higher NK/SKOV3 ratio was observed in the peritoneal lavage for animals treated with gRV-CAR NK cells when *TIGIT* was ablated (**Fig. 5m**).

To determine the role of TIGIT ligands, we characterized their expression in both tumor cell lines. Interestingly, the SKOV3 cell line expressed high levels of CD155 and CD112, the main ligands for TIGIT, whereas their expression was nearly absent on NALM6 cells (**Extended Data Fig. 6c**). Importantly, DNAM-1, an activating NK-cell receptor, competes with TIGIT for these ligands. To investigate the individual and combinatorial effects of these ligands on NK cells, we engineered CD19^+^ SKOV3 cells with deletion of CD155, CD112, or both (double negative, DN) (**Supplementary Fig. 9a**). We designed an *in vitro* competitive cytotoxicity assay where a 1:1 mix of two ligand variants of the CD19^+^ SKOV3 cell line were co-cultured with NK or CAR NK cells in which TIGIT, DNAM-1, or both were ablated (**Supplementary Fig. 10a** and **Supplementary Fig. 9a,b,c**). Target cells expressing CD155 were preferentially depleted by NK cells when DNAM-1 expression was maintained (*TIGIT* KO and *AAVS1* KO conditions), whereas CD112-expressing target cells were preferentially spared when DNAM-1 expression on NK cells was lost (*CD226* KO and *CD226* KO/*TIGIT* KO conditions). However, expression of a CAR by NK cells largely dampened this preferential killing, although *TIGIT* deletion continued to favor the depletion of CD155^+^ targets (**Supplementary Fig. 10b**). The inhibitory effect of *TIGIT* in this short-term co-culture assay was more subtle, but normalization by the double-receptor KO (*TIGIT* KO/*CD226* KO) conditions revealed a significant inhibitory effect when target cells were expressing both the CD112 and the CD155 ligands (**Supplementary Fig. 10c,d**). Overall, the *in vivo* benefit of *TIGIT* KO in NK and CAR NK cells was dependent on the tumor model, and our data suggest the relevance of the expression of the cognate ligands CD155 and CD112 by the tumor cells for enhanced anti-tumor activity.

## Discussion

Modern genome engineering technologies offer powerful opportunities to enhance the anti-tumor activity of NK-cell therapies^49–52^. We present a feeder-free workflow for genetic modification of primary human NK cells that combines CRISPR/Cas9 and recombinant AAV6 for transgene KI. This workflow builds upon a growing corpus of published reports^27,28,30,53,54^ and is compatible with multiple CRISPR-based tools for efficient gene deletion, base editing, and, critically, site-specific transgene integration. It also is compatible with retroviral transduction, providing broad flexibility for advanced NK cell engineering. We believe that this comprehensive toolbox could not only help enhance NK-cell therapies but also serve as a valuable resource to further explore NK-cell biology.

This workflow uses electroporation to deliver the CRISPR/Cas9 components. While electroporation is highly efficient, it is associated with residual toxicity in primary NK cells, despite extensive optimization by our group and others^27,30,55,56^. To address this limitation, alternative delivery strategies could be explored in combination with an AAV6-mediated HDRT. Promising approaches include amphiphilic cell-penetrating peptides, either complexed or covalently linked to RNP complexes^57,58^, gentle mechanical methods adapted from primary T cells^59^ and successive generations of virus-like particles^60–64^ that can be further optimized for NK cells^65^. Lipid nanoparticles also have recently emerged as a vehicle for CRISPR/Cas9 delivery in primary immune cells^66–68^.

A key advantage of KI strategies over semi-random integration methods, such as those mediated by retroviruses or transposases, relates to safety. Precise genomic targeting via guide RNA-directed CRISPR editing reduces the risk of insertional mutagenesis^69^ and enables fine control over integrated transgene copy number (mono- or bi-allelic integration, with a maximum of two copies in a single cell)^70–73^. An additional benefit of this approach results from a more homogenous and predictable transgene expression across edited cells contributing to a consistent and reproducible final cell product^23^. Furthermore, for experimental comparisons between CAR constructs, the calibrated expression levels achieved through targeted integration help mitigate expression bias, a strategy we applied here to the evaluation of CAR ITAM variants.

To our knowledge, this article presents the largest reported series of genomic loci evaluated for transgene integration in primary human NK cells^22,27,28,53,54,56,74–79^, applied here to deliver a CAR transgene. The same approach could be extended to other NK-relevant payloads, such as cytokines, synthetic, optimized or orthogonal receptors, or intracellular signaling adaptors and modulators. In addition, we describe different strategies for modulating CAR expression, which can be a valuable resource for designing future NK therapies. As we previously reported in cell lines and primary human T cells^23,41^, our results demonstrate that both the integration locus and promoter choice can be used to tune transgene expression to a desired level. In another study exploring *CD38* as an integration locus^28^, the EF1α promoter was compared to the endogenous *CD38* promoter, and higher transgene expression was observed with EF1α. Given the packaging size constraints of AAV vectors, using an endogenous promoter, when appropriate, can increase the available cargo space for the transgene.

For certain genomic loci that we tested, such as *FCGR3A*, *NCR1* and *LILRB1*, editing efficiency was relatively low, yet transgene expression was high. In the context of iPSC-derived NK cells, where a master cell bank of a desired single-cell clone is established, editing efficiency is less critical than the expression level achieved in successfully edited cells^74,80^. For preclinical or clinical applications using other NK cell sources, selection strategies can be implemented to enrich for CAR-positive cells, if needed, as reported in previous studies^70,81–83^.

Blocking or ablating NK inhibitory receptors or negative regulators has long been explored as a potential strategy to enhance the anti-tumor activity of NK or CAR NK cells^24,25,50,84^. Here, we propose a single-step editing approach that couples CAR KI and endogenous receptor KO. We successfully applied this strategy to four inhibitory receptors (*TIGIT*, *KLRB1*, *KLRC1*, *LILRB1*) that each have been reported as promising targets for genetic ablation^56,85–100^.

An important observation from our study is the lack of direct correlation between *in vitro* and *in vivo* results. This result is exemplified by *KLRC1*-CAR NK cells (our top-performing candidate *in vitro*) that failed to exhibit superior activity *in vivo* against the SKOV3 model. A similar discrepancy was observed when comparing CAR ITAM variants, highlighting the importance of early *in vivo* evaluation of therapeutic candidates in appropriate preclinical models. CRISPR-mediated deletion of *KLRC1* previously has been shown to enhance *in vivo* anti-tumor efficacy of anti-CD33 CAR NK cells against acute myeloid leukemia^86^. Therefore, while targeting a CAR to the *KLRC1* locus was ineffective in the SKOV3 model, this strategy may still hold therapeutic potential in other settings^56,86,99,100^.

More broadly, our results support the notion that there may be no universally optimal locus for CAR integration in primary NK cells. Instead, the ideal integration site (and potential gene deletion) might depend on the tumor type and therapeutic context. As a proof of concept, we screened our four top candidates (selected based on CAR expression) in an *in vivo* ovarian cancer xenograft model, which identified *TIGIT* as a promising integration platform in this setting. Other groups have investigated *TIGIT* deletion in CAR NK cells across *in vitro* and *in vivo* tumor models with inconsistent results^65,70,89,101^. Similarly, our study highlights model-dependent differences in the functional impact of *TIGIT* deletion, which may reflect variable expression of its cognate ligands as well as other factors such as the complex interactions with its paired receptors DNAM-1, CD96, and PVRIG^102,103^.

In summary, our study provides a versatile genome engineering approach for primary human NK cells that enables efficient site-specific transgene integration with tunable expression and that allows for concurrent deletion of a negative regulator of NK cell activity. This approach lays the groundwork for next-generation NK cell therapies designed to harness NK-cell biology where multiple synergistic and precise genetic edits may be essential to overcome intrinsic limitations of NK cells and address the challenges posed by the complex environments of hematologic malignancies and solid tumors.

## Methods

### Primary NK cell isolation and expansion

Methods for expansion and genome editing of primary human NK cells were adapted from a previously described workflow^27^. Primary human NK cells were isolated from PBMCs obtained after density gradient centrifugation of Trima residuals from apheresis collection (Vitalant). Negative selection of NK cells was performed using an EasySep™ Human NK Cell Enrichment Kit (Stemcell) according to the manufacturer instruction. Cells were cultured in NK MACS medium (Miltenyi) supplemented with 5% human platelet lysate (HPL) (EliteGro-Adv, Elite Cell), 0.5% penicillin-streptomycin (Gibco) and 1000 U/mL recombinant human IL-2 (Peprotech). Cells were activated over 7 days with anti-CD2-coated and anti-NKp46-coated beads (Miltenyi) at a cell:bead ratio of 2:1 and cultured at 1 x 10^6^ cells/mL in 24-well plates incubated at 37° C 5% CO_2_. Cells were left undisturbed until day 5. At day 5, half of the medium was exchanged with fresh medium and IL-2. At day 7, the cells were collected and activation beads were removed using an EasySep™ Magnet. Starting at day 7, cells were replated twice per week at 1 x 10^6^ cells/mL with fresh medium and 1000 U/mL IL-2 for further expansion. After bead removal, 24-well GREX plates (Wilson Wolf Manufacturing) could also be used for larger-scale production. Each well was filled with 8 mL of complete medium. Twice per week, 6 mL medium was removed and replaced with fresh complete medium and fresh IL-2 (final concentration of 1000 U/mL). When the cell number reached 50 x 10^6^ cells in a well, cells were split and reseeded at ∼10 x 10^6^ cells per well for further expansion.

### NK cell editing

Two million NK cells were electroporated with 40 pmol of purified Cas9-NLS protein (MacroLab) complexed with 80 pmol of sgRNA (Synthego) using a 4D nucleofector (Lonza) in 20 μL of P3 buffer and supplement (Lonza) using the CM-137 pulse code. Immediately after electroporation, edited NK cells were rescued in pre-warmed culture medium and plated for cell culture. For targeted integration, NK cells were resuspended at 3 x 10^6^ cells/mL in complete NK medium after electroporation, assuming a loss of 50% of the cells. After a recovery time of 30-60 minutes, NK cells were transduced with AAV6 using a multiplicity of infection (MOI) ranging from 1 x 10^4^ to 1 x 10^5^ vg/cell. The DNA-PK inhibitor M3814 (Nedisertib, 0.5 μM, ChemieTek) was added in the same step as transduction in certain experiments to enhance KI efficiency. After an overnight incubation, edited cells were resuspended in fresh complete medium at 1 x 10^6^ cells/mL for further expansion. For large-scale production in the context of *in vivo* experiments, electroporated NK cells were directly plated in 24-well GREX plate at 5 x 10^6^ cells/mL (assuming a loss of 50% of the cells after electroporation), with a minimum of 3 x 10^6^ cells seeded per well (0.6 mL). AAV6 and M3814 were added in the well for transduction. After an overnight incubation, prewarmed media was directly added into the GREX well for a minimum dilution of 5-fold without media exchange. The quantity of medium was increased over the course of 3 days to reach the final volume of 8 mL per well. For base editing experiments, 2 µg mRNA encoding an ABE8e(V106W) adenine base editor produced by *in vitro* transcription were mixed with 1.5 µg sgRNA (Synthego) and added per 2 x 10^6^ NK cells resuspended in 20 µL of P3 buffer with supplement, as previously reported with T cells^32^. Electroporation and cell recovery was similar to the strategy described for NK cell gene ablation.

### sgRNA design and efficiency assessment

*In silico* tools (www.benchling.com and https://design.synthego.com) were used to design sgRNA sequences. sgRNAs were validated by evaluating the KO efficiency by flow cytometry usually four or five days after editing. KO efficiency was also evaluated at the genomic level using amplicons generated by PCR around the anticipated cut site. Amplicons were submitted for Sanger sequencing and analyzed by Inference of CRISPR Edits (ICE) for estimating a KO score and an indel score (https://ice.synthego.com). For base editing assessment, amplicons were sequenced using the Nanopore-based Premium PCR service (Plasmidsaurus), and raw sequencing reads were aligned against the linear consensus sequence (Plasmidsaurus). The sgRNAs and corresponding primers for ICE analysis are listed in **Supplementary Table 1**.

### AAV production

AAV production was carried out as previously described^104^. Briefly, AAV2-ITR-containing plasmids were used to package transgenes of interest into AAV6 capsids by transfection of HEK293T cells together with adenovirus helper and AAV Rep-Cap plasmids using polyethylenimine. Packaging cells were collected and lysed 3 d after transfection. After treatment with Benzonase (Millipore Sigma), AAV vectors were purified using iodixanol gradient ultracentrifugation (OptiPrep, StemCell Technologies). AAV extracts were collected and the vectors were concentrated while iodixanol was washed by several centrifugation cycles with a 50-kDa Amicon column (Millipore Sigma). Purified AAV vectors were stored at -80° C. AAV vector titers were determined by qPCR on DNase I (NEB)-treated, proteinase K (Qiagen)-digested AAV samples, using primers targeting the viral genome. qPCR was performed with SsoFast Eva Green Supermix (Bio-Rad) on a StepOnePlus Real-Time PCR System (Applied Biosystems). Relative quantity was estimated by comparison to a serial dilution of vector plasmid standards of known concentrations.

### NK cell transduction by retrovirus

Activated NK cells were transduced by spinfection at day 7. Briefly, debeaded NK cells resuspended in fresh NK medium and 1000 U/mL IL-2 were mixed with a 20-fold concentrated gammaretroviral (gRV) vector encoding a transgene of interest for a final concentration of 2×10^6^ cells/mL. The usual volume ratio for mixing cells and retrovirus was 2:1. The cell mixture was distributed in washed Retronectin (Takara)-coated (20 µg/mL in sterile PBS overnight) non-TC treated plates (1×10^6^ cells per well for a 24-well plate). Plates were centrifugated for 60-90 min at 2000 x g (30° C) and then placed in the incubator overnight. The next day, cells were collected, spun, and resuspended in fresh medium at 1×10^6^ cells/mL for further cell culture.

### RD114-pseudotyped gammaretrovirus production

Plasmids encoding a SFG gRV vector^42,105^ were cloned to carry a transgene of interest and purified with the NucleoBond Xtra Maxi kit (Macherey-Nagel). Lipofectamine LTX and Plus Reagent (Invitrogen) were used for transfection of 293Vec-RD114 packaging cells^31^ (Biovec Pharma). The supernatant containing the retroviral vectors was collected at 48 and 72 h post-transfection and filtered. Retroviral vectors were concentrated 20-fold with RetroX concentrator (Takara) according to the manufacturer instructions before aliquoting in vials for single use and storage at -80° C.

### Target tumor cell lines

Cell lines were maintained in sterile conditions in a 5% CO_2_ incubator at 37° C. Firefly luciferase^+^ GFP^+^ NALM6 (naturally CD19^+^) cells were cultured in RPMI-1640 (Gibco, 2 mM L-glutamine) supplemented with FBS (Corning, 10%), sodium pyruvate (Life Technologies, 1%), HEPES buffer (Corning, 1%), penicillin– streptomycin (Gibco, 1%), non-essential amino acids (Gibco, 1%), and 2-mercaptoethanol (Gibco, 0.1%). SupB15 cells (ATCC, naturally CD19^+^) were transduced with a lentiviral vector to express firefly luciferase, mKate2, and a blasticidin resistance gene, and polyclonal transduced cells were selected using blasticidin and subsequently sorted by flow cytometry. Blasticidin was maintained in the culture medium. The cells were cultured in IMDM (Gibco) supplemented with FBS (20%), 2-mercaptoethanol (0.08%), and penicillin-streptomycin (1%). Clonally selected firefly luciferase^+^ mKate2^+^ NGFR^+^ SKOV3 cells were transduced with a lentiviral vector for expressing human CD19 and a puromycin resistance gene, and cells were selected with puromycin. Puromycin was subsequently maintained in the culture medium. hCD19^+^ SKOV3 cells and the hCD19^-^ parental cell line were cultured in high-glucose DMEM with GlutaMAX (Gibco) supplemented with FBS (10%), HEPES buffer (1%), sodium pyruvate (1%), and penicillin-streptomycin (1%). For CD155 and CD112 KO variants, deletion of *PVR* and/or *NECTIN2* was performed using CRISPR/Cas9 followed by magnetic-based negative selection using LS and LD columns (Miltenyi). RFP^+^ CD19^+^ A375 cells (kind gift from Franziska Blaeschke) were cultured in high-glucose DMEM with GlutaMAX supplemented with FBS (10%), HEPES buffer (1%), sodium pyruvate (1%), and penicillin-streptomycin (1%). Clonally selected firefly luciferase^+^ mKate2^+^ NGFR^+^ K562 cells and firefly luciferase^+^ blasticidin resistant RAJI cells (ATCC) were cultured in RPMI-1640 (2 mM L-glutamine) supplemented with FBS (10%) and penicillin-streptomycin (1%).

### Packaging cell lines

HEK293T cells (ATCC) and 293Vec-RD114 cells (Biovec Pharma) were cultured in high-glucose DMEM with GlutaMAX supplemented with FBS (10%), sodium pyruvate (1%), HEPES buffer (1%), and penicillin– streptomycin (1%). HEK293T cells were used for AAV6 packaging and 293Vec-RD114 cells were used for retrovirus packaging.

### Flow cytometry

Flow cytometry was performed on an LSR Fortessa X-50 flow cytometer (BD Biosciences). Cells were resuspended in FACS buffer (phosphate-buffered saline (PBS), 2% FBS, and 1 mM EDTA) and stained with live–dead stain (15 min, room temperature) before applying antibodies targeting surface markers (25 min, 4° C). Zombie NIR (BioLegend) and Ghost Dye Red 780 (Tonbo Biosciences) were used for live–dead staining. Human Fc Receptor Blocking Solution (BioLegend) and Brilliant Stain Buffer Plus (BD Biosciences) were used as needed. Cytometry data were processed and analyzed using FlowJo software (BD Biosciences).

Spectral flow cytometry was performed on a five-laser Cytek Aurora Spectral Cytometer. Cells were washed in FACS buffer and treated with Human Fc Receptor Blocking Solution (BioLegend) and True-Stain Monocyte Blocker (BioLegend) while stained with viability dye for 25 min at 4° C. Cells were then stained with antibodies targeting surface markers for 30 minutes at 4° C. Cells were permeabilized and fixed using FluoroFix (BioLegend). Antibodies targeting intracellular markers were added, and cells were stained for 30 min at 4° C. Brilliant Stain Buffer Plus was used as necessary. Unmixing errors were corrected by spillover correction, and all analysis was processed using FlowJo software.

A list of the antibodies used is reported in **Supplementary Table 2**. For CAR staining, two different reagents were used. In some experiments, cells were stained with polyclonal Alexa Fluor 647-conjugated goat anti-mouse F(ab’)_2_ (Jackson ImmunoResearch) and then blocked with normal mouse serum (Millipore Sigma) before further antibody staining was performed. In other experiments, a labelled antibody targeting the Glycine-Serine linker of the CAR was used (G4S, Cell Signaling). For quantification of cells during flow cytometry, 30 μL of CountBright Absolute Counting Beads (Invitrogen) were added to each sample following the manufacturer’s protocol. For phospho-flow staining, stimulated cells were fixed by adding an equivalent volume of prewarmed Cytofix solution (BD Biosciences) and incubated at 37° C for 15 min. Cells were washed and permeabilized using prechilled Perm Buffer III (BD Biosciences) for 30 min on ice. Cells were washed for subsequent antibody staining. For degranulation and cytokine secretion assays, NK cells were stimulated for 6 h by target cells at a defined effector-to-target ratio, in presence of anti-CD107a antibody. After 1 h, monensin (1000x stock, BioLegend) was added to the co-culture. The stimulated cells were collected and then fixed after viability and extracellular staining using 100 μL of CytoFast Fix Perm solution (BioLegend) for 20 min. Cells were subsequently permeabilized using CytoFast Perm Wash solution (BioLegend) according to the manufacturer’s instructions, before intracellular staining for 30 min on ice.

### Luciferase-based cytotoxicity assays

The cytotoxicity of NK cells was determined by using a standard luciferase-based assay^23^. In brief, several cell lines were transduced with lentivirus to express firefly luciferase and served as target cells. The effector (E) and tumor target (T) cells were co-cultured in triplicate at the indicated E:T ratios using white-walled 96-well flat clear-bottom plates with a given number of target cells in a total volume of 100 μL per well in NK cell medium without IL-2. The control for *maximal* signal was the target cells alone, and the control for *minimum* signal was the target cells with Tween-20 (0.2%). Co-cultures were incubated for approximately 20-24 h. Then, 100 μL D-luciferin (GoldBio, 0.75 mg/mL) was added to each well, and luminescent signal was measured using a GloMAX Explorer microplate reader (Promega). Cytotoxicity was calculated as follows: 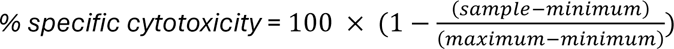. The number of target cells per well was 4-5 x 10^4^ target cells for cells in suspension (NALM6, SUPBB15) and 2 x 10^4^ target cells seeded a day before for adherent cells (SKOV3).

### Incucyte cytotoxicity assays

NK cells were co-cultured with 0.5-1×10^4^ pre-plated RFP^+^ A375 cells in complete NK medium in 96-well flat bottom plates. Each condition was plated in triplicate. The tumor cell count per well was quantified every 4 h over a 7-8 day period using IncuCyte S3 live-cell imaging (Sartorius). It was calculated as the number of objects exhibiting red fluorescence at a given time point, normalized to the number of objects exhibiting red fluorescence at the start of the assay. Tumor-only wells were used as positive controls.

### Competitive cytotoxicity assay

CD19^+^ SKOV3 cells with specific deletion of CD155, CD112, or both, and CD155^+^ CD112^+^ parental cells were used as targets for an NK cell competitive cytotoxicity assay. Target cells of interest were labelled in prewarmed PBS using CellTrace Blue (CTB) or CellTrace Violet (CTV) (Invitrogen) according to manufacturer’s instructions. Cells were then collected, counted, and resuspended in complete NK cell medium. Distinct mixes (1:1 ratio) were prepared with differentially labelled (CTB or CTV) target cells. For each target mix, a mirror mix was also prepared by inverting which cell variants were labelled with CTV and CTB. Mixed target cells were seeded in a flat-bottom 96-well plate (4 x 10^4^ total cells per well) and incubated undisturbed for 4 h to allow adherence. NK and gRV-CAR^+^ NK cells deleted for *CD226*, *TIGIT,* or both were used as effectors, with *AAVS1* KO NK cells used as DNAM1^+^ TIGIT^+^ NK controls. NK cells were then carefully added to the target cells at a 1:1 effector to target (E:T) ratio and cocultures were incubated for approximately 20 h. Each condition was plated in duplicate. At the end of the assay, adherent cells were detached using TrypLE (Gibco), transferred in a V-shape plate for viability staining and stained for surface markers in cold FACS buffer allowing to distinguish target cells from remaining NK cells. Cells were fixed using Cytofix solution (BD Biosciences) for flow cytometry analysis. For each well, the relative enrichment of CTB^+^ versus CTV^+^ surviving target cells was assessed by calculating the CTV^+^/CTB^+^ ratio. Mirror mixes data were pooled together to obtain final quadruplicate data for each coculture condition. Normalization was done using the CTV^+^/CTB^+^ ratio obtained for the same target mix with the indicated control effector condition.

### *In vivo* mouse models

NOD.Cg-*Prkdc^scid^ Il2rg^tm1Wjl^*/SzJ (NSG) mice bred at the University of California, San Francisco (UCSF) or purchased from The Jackson Laboratory were handled in accordance with a protocol approved by the UCSF Institutional Animal Care and Use Committee (IACUC). Before and during the experiment, mice were kept on Clavamox antibiotic. For the hCD19^+^ SKOV3 model, female NSG mice were injected with 0.1 x 10^6^ ffLuc^+^ hCD19^+^ SKOV3 (intraperitoneally - i.p.) 4 to 8 days before the i.p. injection of 5-7 x 10^6^ (CAR) NK cells. For some experiments, mice were injected i.p. twice weekly with IL-2 (50000 U) for 3 to 9 weeks. For the NALM6 model, 8 to 12 weeks old male NSG mice were injected with 0.1 x 10^6^ ffLuc^+^ NALM6 (i.v.) 4 days before the i.v. injection of 10 x 10^6^ (CAR) NK cells. BLI was performed the day before NK injection and then one or two times per week using a Xenogen *in vivo* imaging system. After the first BLI measurement (one day before NK cell injection), mice were assigned to each treatment condition to keep a similar average mass, age, and tumor burden across conditions. At each imaging session, mice were injected i.p. with luciferin (3 mg luciferin in 0.2 mL DPBS) and anesthetized with isoflurane (Medline Industries). The default imaging time was 1 min, and shorter exposures were used for images that had a saturating signal. Luminescence was quantified using Living Image software (PerkinElmer). Reported BLI radiance values are an average from imaging mice on their front and back. Mice were euthanized according to the approved protocol if they reached morbidity end points. For certain experiments, an additional endpoint based on tumor burden was included (mean average radiance of 2 x 10^7^ p.s^-1^.cm^-2^. Sr^-1^ for **Fig. 3l** and 5 x 10^6^ p.s^-1^.cm^-2^. Sr^-1^ for **Fig. 4h**).

### Peritoneal lavage and spleen analysis

Mice were euthanized and the peritoneal cavity was washed twice with 5 mL ice-cold PBS with 1 mM EDTA using a 21G ¾ needle with tubulure (Vacutainer). After abdominal massage, the washing liquid was carefully collected and strained using 70 µm cell strainers (Corning). Cells were treated for 2 min with 1 mL ACK lysing buffer (Quality Biological) for red blood cell lysis. After quenching with FACS buffer, remaining cells were resuspended in a defined volume of FACS buffer and stained for flow cytometry using mouse FcR blocking reagent (Miltenyi) to avoid unspecific labelling. Spleens were harvested following the peritoneal lavage collection, crushed, and strained before ACK lysing buffer treatment for 2 min. After quenching, remaining cells were strained again and resuspended in a defined volume for flow cytometry staining. Mouse FcR blocking reagent was used before labelling.

### Graphical and statistical analysis

Data were analyzed using Prism version 10.3.1 (GraphPad) and R version 4.2.1 (R Core Team, https://www.R-project.org) with the packages ggplot2, ggpubr, survival, survminer through the RStudio version 2022.07.2 software. Data are expressed as mean ± SEM or median ± inter-quartile range (IQR). Repeated-measures one-way ANOVA, paired *t*-tests, Mann-Whitney, Kruskal-Wallis and log-rank tests were used as indicated in each figure legend, using Tukey’s and Dunn’s tests for multiple comparison correction. Where applicable, all statistical tests were two-tailed with a significance level of 0.05 and reported as: *, *p* ≤ 0.05; **, *p* ≤ 0.01; ***, *p* ≤ 0.001; ****, *p* ≤ 0.0001.

## Acknowledgements

This work has been supported by the Fondation ARC pour la recherche sur le cancer www.fondation-arc.org. The UCSF Parnassus Flow Core RRID:SCR_018206 is supported by the DRC Center Grant NIH P30 DK063720. C.R.C. was supported by the UCSF Medical Scientist Training Program (T32GM141323). C.C.W. was supported by an NCI F99/K00 Fellowship (K00CA245718). Figures were produced using elements from https://www.biorender.com. We thank all current and former members of the Eyquem, Lanier, and Aguilar labs, as well as Sophie Caillat-Zucman, for their valuable input and support during this project.

## Author contributions

V.A. and J.E. conceived the study. V.A. and A.G.R. designed and performed experiments and analyzed data. W.A.N., J.J.M., J-Y.C., An.T., G.R.K., C.L., R.S. and C.C.W. conceived and prepared reagents. P.L.B., Al.T., T.T., Y.M., J.S., Z.L., A.S. and O.A.A. performed experiments. P.L.B., W.A.N., J.J.M, C.R.C., A.S., A.M., L.L.L. and O.A.A. contributed to experimental design. V.A., A.G.R., L.L.L., O.A.A. and J.E. wrote and edited the manuscript. All authors read the manuscript and agree with its contents.

## Data Availability

All data are available from the corresponding authors on reasonable request.

## Extended Data Figures

**Extended Data Figure 1.**
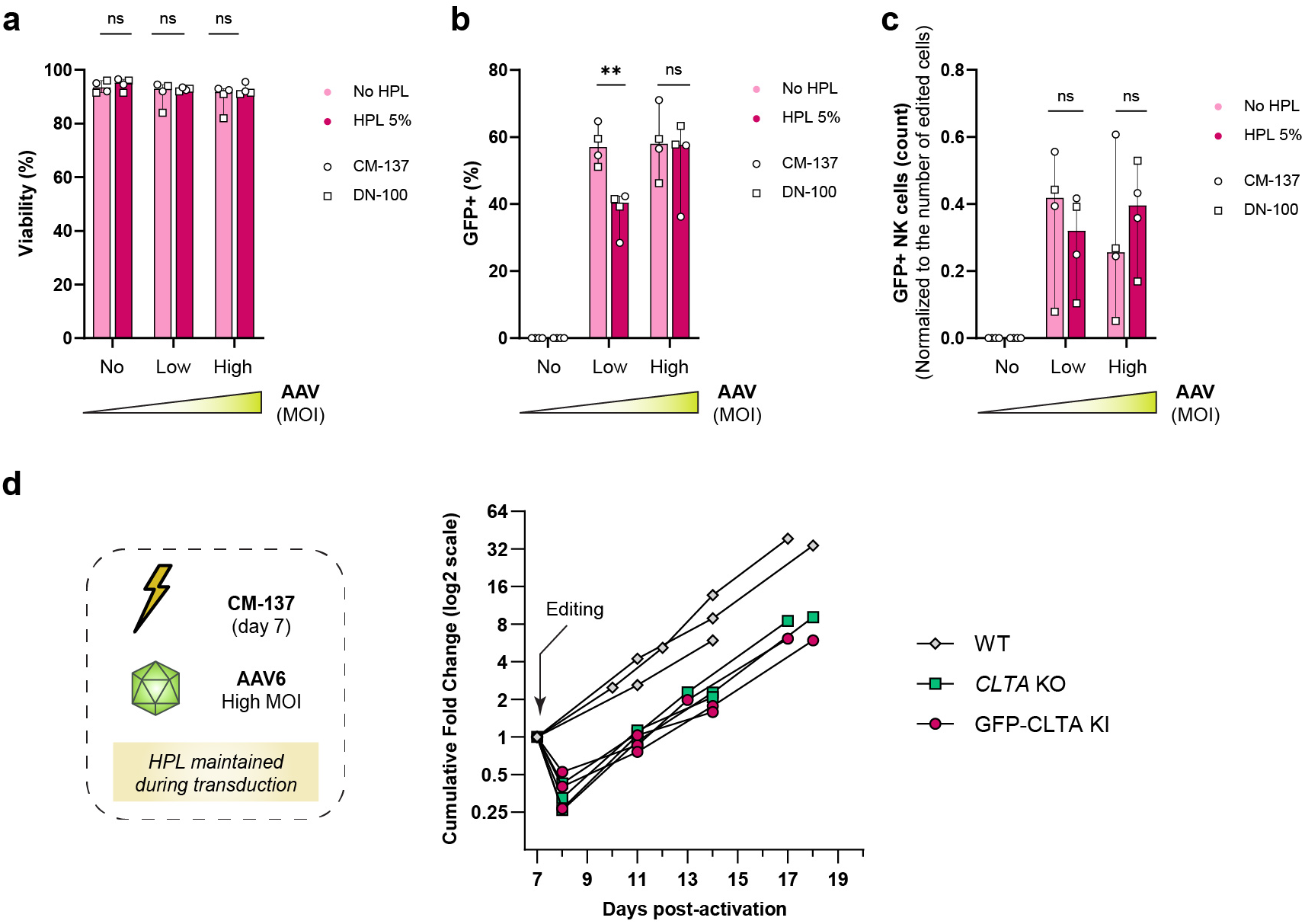
Influence of HPL during AAV transduction on editing outcomes (related to Fig. 1) **a-c**, Targeted integration of GFP at the *CLTA* locus was performed on primary NK cells, without (light pink) or with (dark pink) HPL during AAV6 transduction. Different MOI were used (low MOI is 1×10^4^; high MOI is 3-5×10^4^). The electroporation code (DN-100 or CM-137) is indicated. Outcomes were assessed 4 d after electroporation: viability (**a**), percentage of GFP+ cells (**b**) and overall editing yield (**c**). Data are shown as median ± range. *P*-values are from a two-tailed paired *t*-test. *n*=3 different donors in 3 independent experiments. **, *p* ≤ 0.01; ns, not significant. **d**, Cumulative fold-change from d 7 (editing day) for the total NK cell number from 3 independent donors, edited with the final optimized conditions (CM-137 electroporation pulse code, 3×10^4^ “high” AAV6 MOI, HPL during AAV6 transduction). A logarithmic scale is used for the y-axis.

**Extended Data Figure 2.**
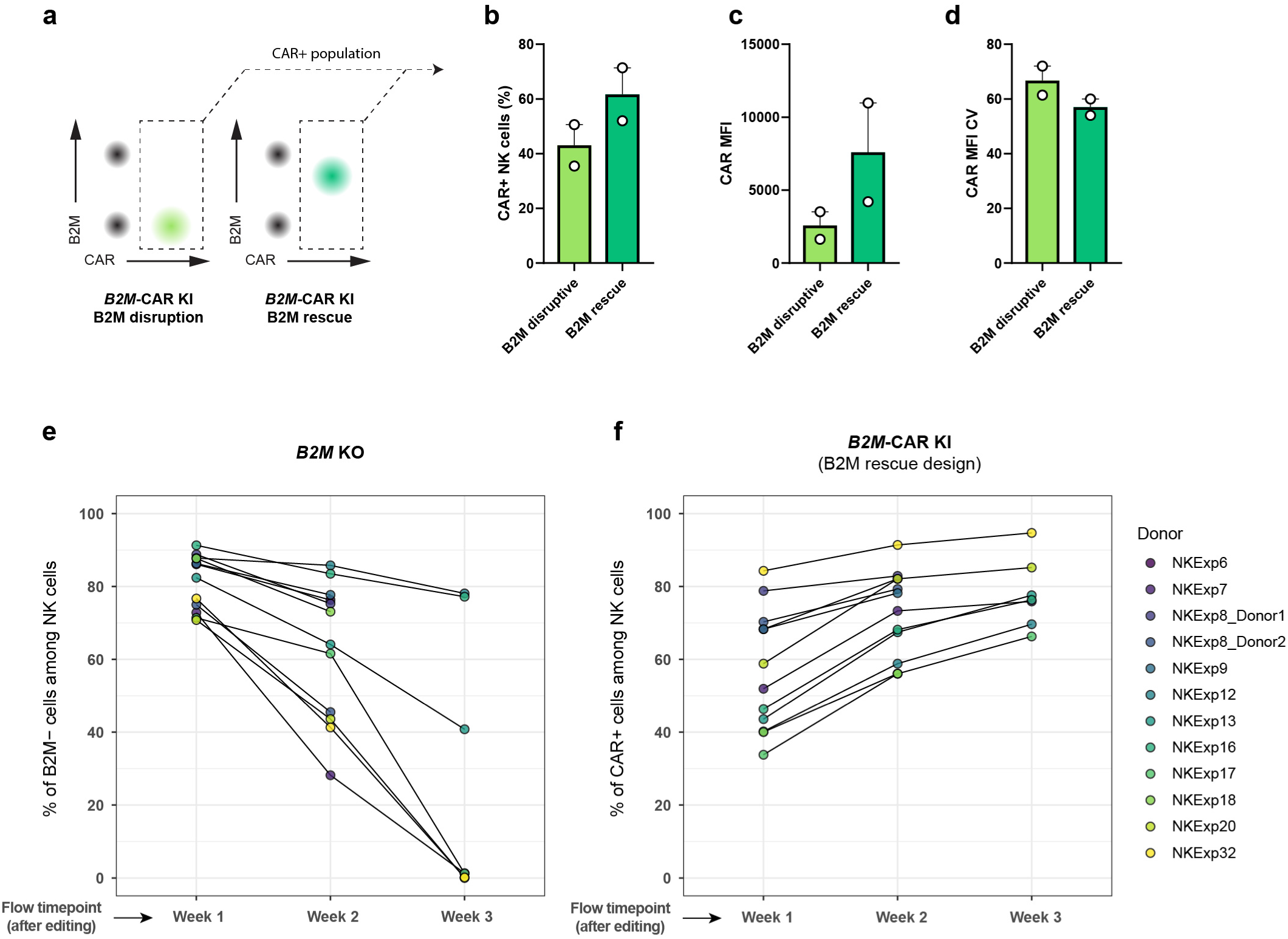
*B2M* rescue design protects *B2M*-CAR NK cells from fratricide despite intermediate B2M expression (related to Fig. 3) **a**, Schematic of the CAR+ NK populations studied in panels (**b-d**) for NK cells edited with disruptive and rescue *B2M-*CAR KI designs. **b**, CAR percentage. **c**, CAR mean fluorescence intensity (MFI) in CAR^+^ NK cells. **d**, CAR MFI coefficient of variance (CV) in CAR^+^ NK cells. Data are shown as mean ± SEM. *n*=2 donors (2 independent experiments). **e-f**, Compilation of data from 12 donors (11 independent experiments) showing the progressive loss of the B2M-negative fraction for NK cells edited for *B2M* KO (**e**), and the progressive enrichment of the CAR^+^ fraction for NK cells edited for *B2M*-CAR KI with a *B2M* rescue design (**f**).

**Extended Data Figure 3.**
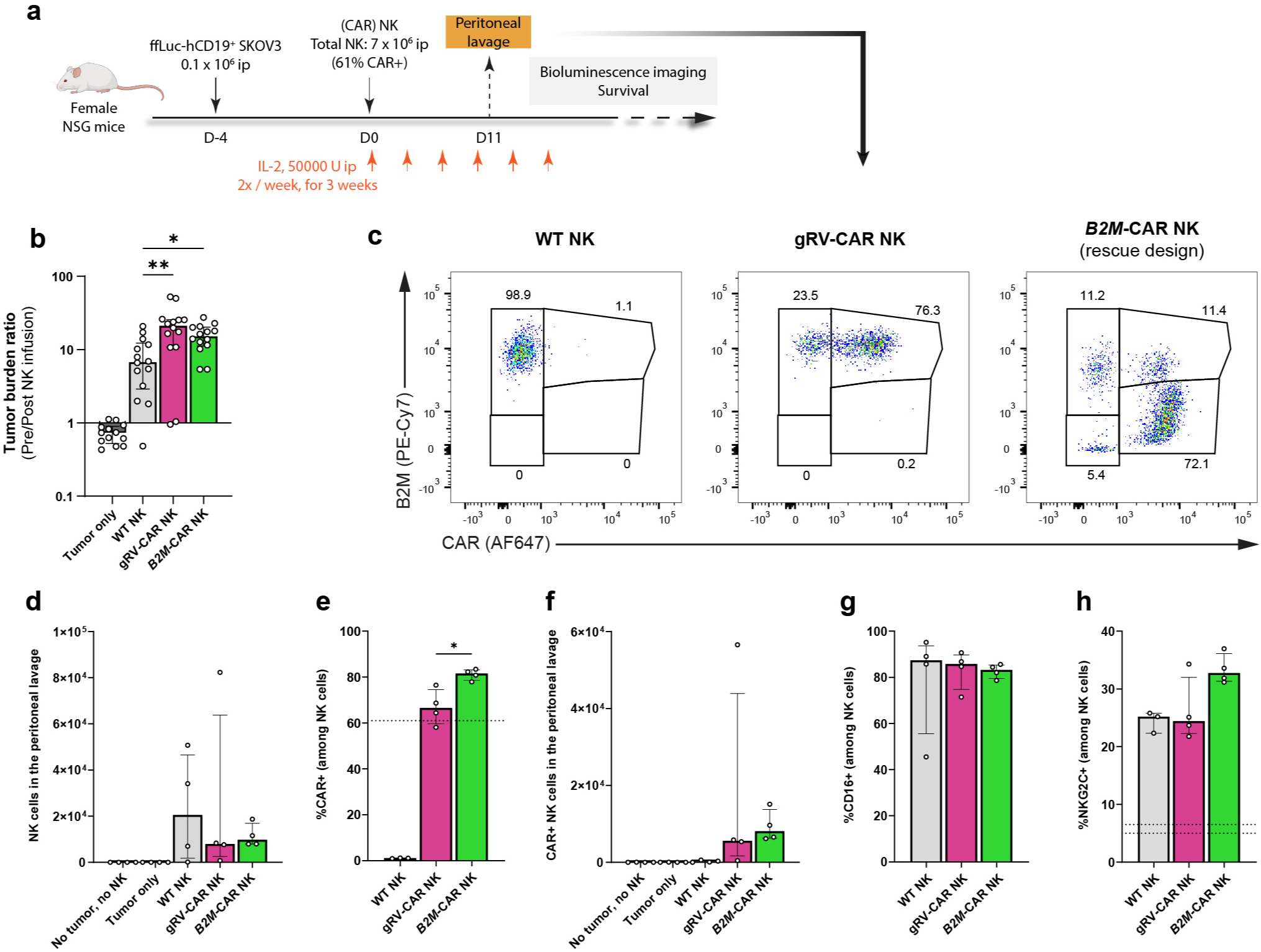
Peritoneal lavage evaluation in a SKOV3 xenograft mouse model (related to Fig. 3) **a**, Schematic experimental design of the SKOV3 xenograft mouse model for *in vivo* assessment of CAR NK anti-tumor efficacy. **b**, Ratio between the tumor burden (mean average radiance assessed by bioluminescence imaging) between day -1 (pre-NK infusion) and during the first week post injection (mean of the tumor burden at day 3 and day 6) highlighting the early response to the treatment. *P*-values are from Kruskal-Wallis test with Dunn’s multiple comparison tests against the WT NK condition. **c**, Representative flow plots (B2M and CAR staining) of the cells recovered from the peritoneal lavage after the euthanasia of a predetermined subset of mice at day 11. The cells are gated on live human CD45^+^ CD3^-^ CD56^+^ cells. **d-h**, Quantification and immunophenotype of the cells recovered from the peritoneal lavage. Number of NK cells recovered from the lavage (**d**), percentage of CAR^+^ cells among NK cells (**e**), number of CAR^+^ NK cells recovered from the lavage (**f**), percentage of CD16^+^ cells among recovered NK cells (**g**), and percentage of NKG2C^+^ cells among NK cells (**h**). For reference, the dotted lines correspond to the baseline CAR percentage in the injected products, and the range of NKG2C percentage before injection. For (**e**), *P*-value is from unpaired two-tailed Mann-Whitney test. Data are shown as median ± IQR. *, *p* ≤ 0.05; **, *p* ≤ 0.01.

**Extended Data Figure 4.**
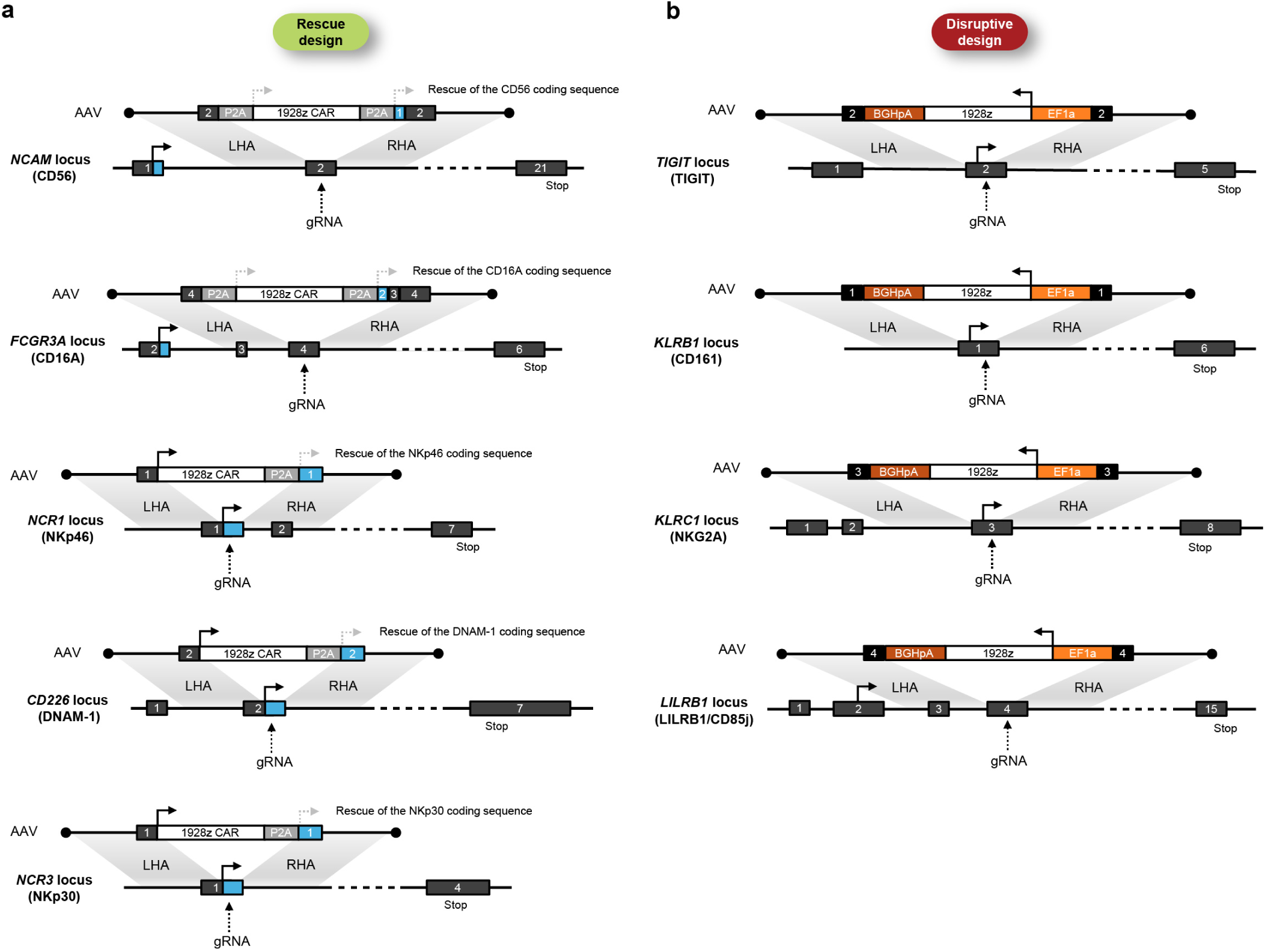
Schematic of the gene editing strategy for CAR integration at activating and inhibitory receptor loci (related to Fig. 4) **a**, A non-disruptive/rescue strategy was designed for activating receptors by integrating the CAR transgene at the beginning of the coding sequence of the targeted locus, followed by a ribosomal skipping P2A peptide allowing to rescue the upstream coding sequence of the targeted gene. The rescue sequence is codon optimized to avoid non-desirable homologous recombination. With this strategy, the CAR is under the control of the endogenous promoter. **b**, A disruptive strategy was designed for inhibitory receptors by integrating the CAR transgene in a reverse orientation compared to the native locus. With this strategy, the CAR is under the control of the EF1α exogenous promoter.

**Extended Data Figure 5.**
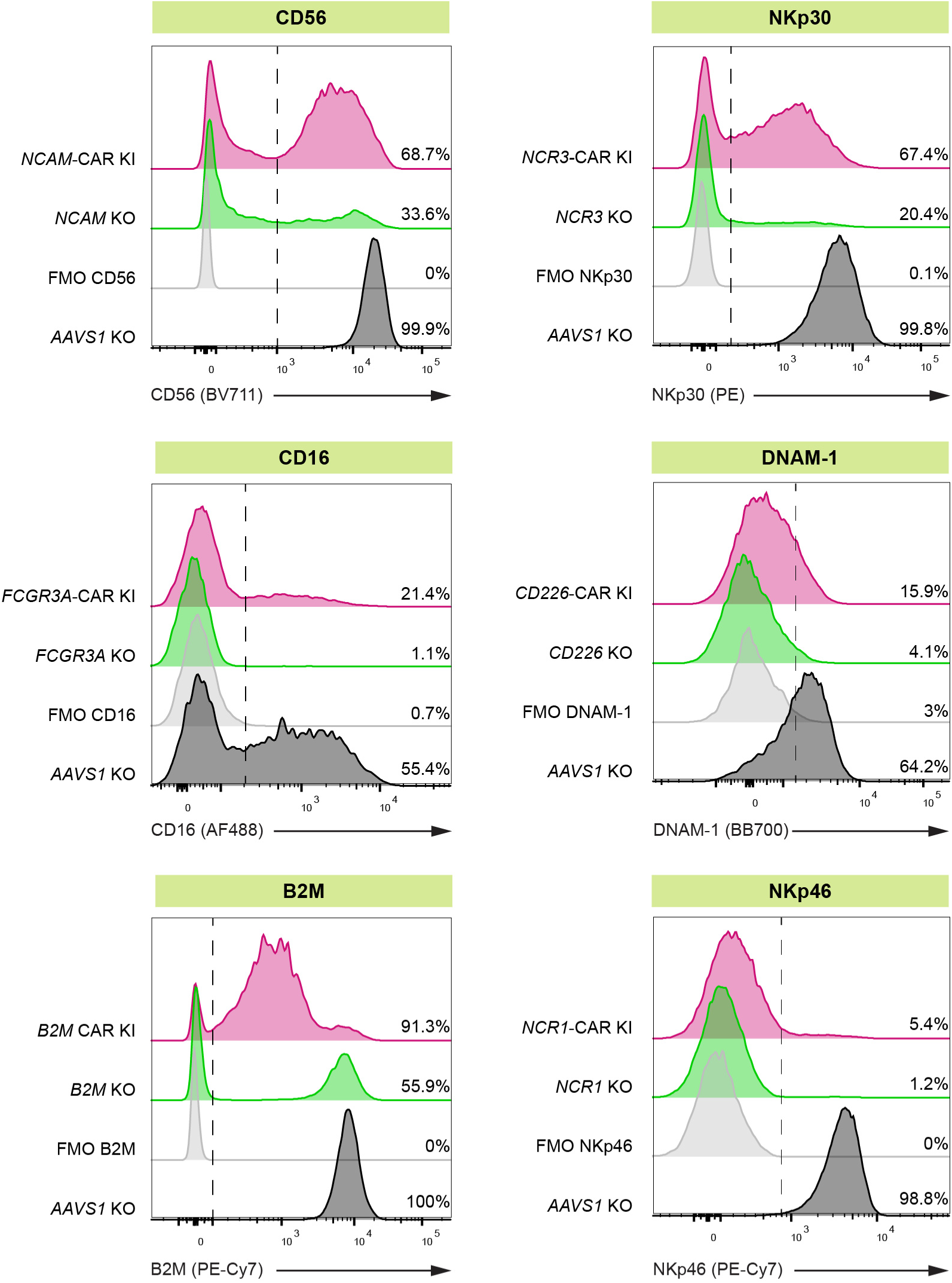
Rescue of the expression of the targeted locus upon CAR knock-in in the context of activating receptors (related to Fig. 4) Targeted integration of a CAR at different NK-activating receptor (rescue design) was performed as described in **Extended Data Fig. 4**. Representative flow cytometry histogram showing the percentage of cells positive for the indicated locus. For each locus, *AAVS1* KO NK cells are used as positive control, and FMO are used as negative control. The receptor expression pattern is shown for both the KO of the given locus and the CAR KI with a rescue design.

**Extended Data Figure 6.**
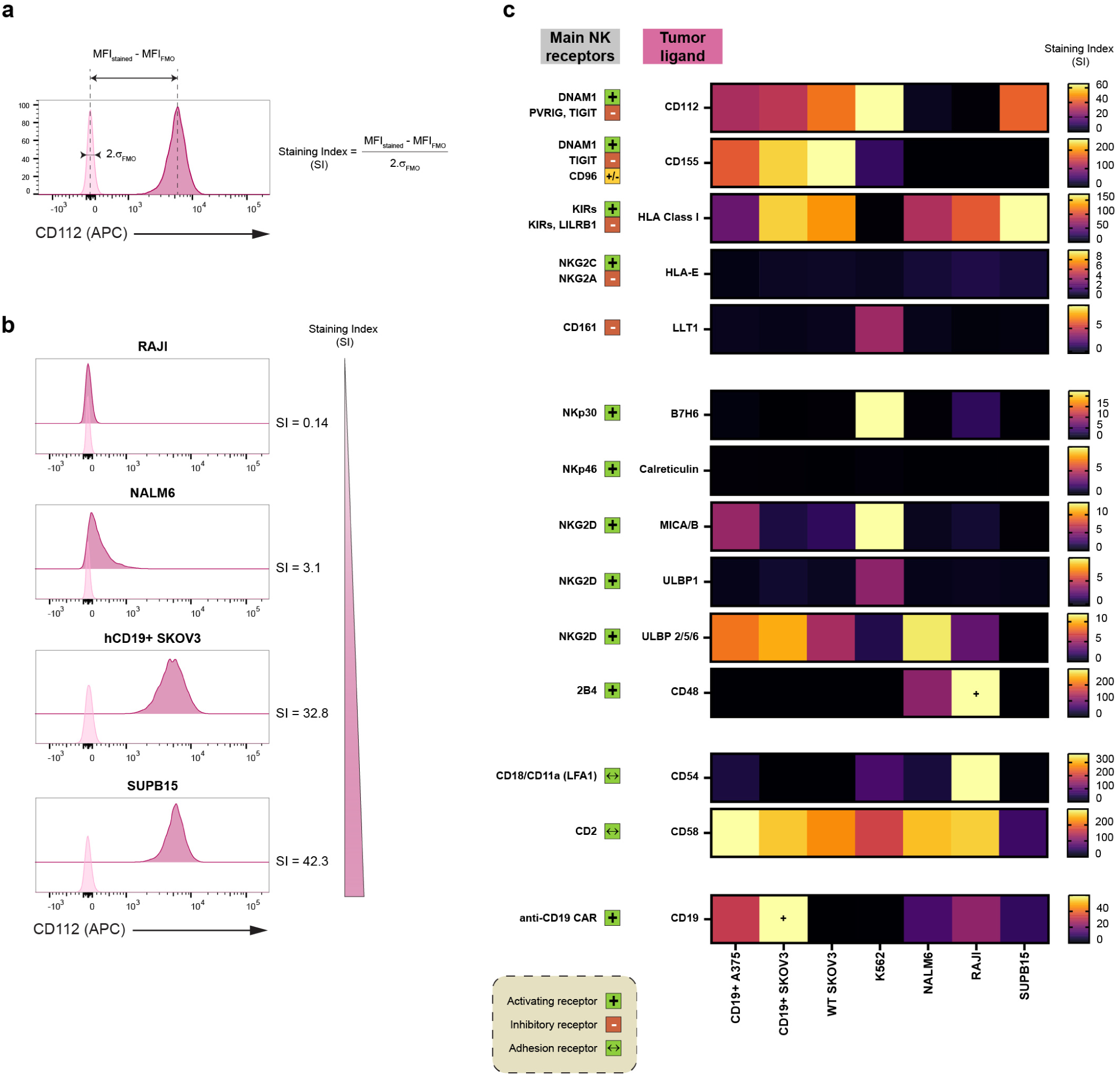
Expression of key NK receptor ligands on the surface of several tumor cell lines. Three multicolor flow cytometry panels were designed using bright fluorophores and compatible with the fluorescent proteins expressed by tumor cell lines (GFP, mKate2), allowing to investigate the expression of the main NK receptor ligands on their surface. **a**, To compare different cell lines while accounting for differences in background noise, a staining index (SI) was used for each ligand. The SI definition was based on the Mean Fluorescence Intensity (MFI) of the sample and the respective FMO used as negative control, and on the standard deviation σ of the FMO. **b**, Example of the staining obtained for CD112 for different cell lines and the corresponding Staining Index. **c**, Heatmap representing the expression (Staining Index) for different NK ligands. Each value represents the mean of 2 technical replicates. The main NK receptors known to bind the corresponding tumor ligands are indicated on the left. The + indicates when there is saturation of the staining index based on the color scale used for the heatmap.

**Extended Data Figure 7.**
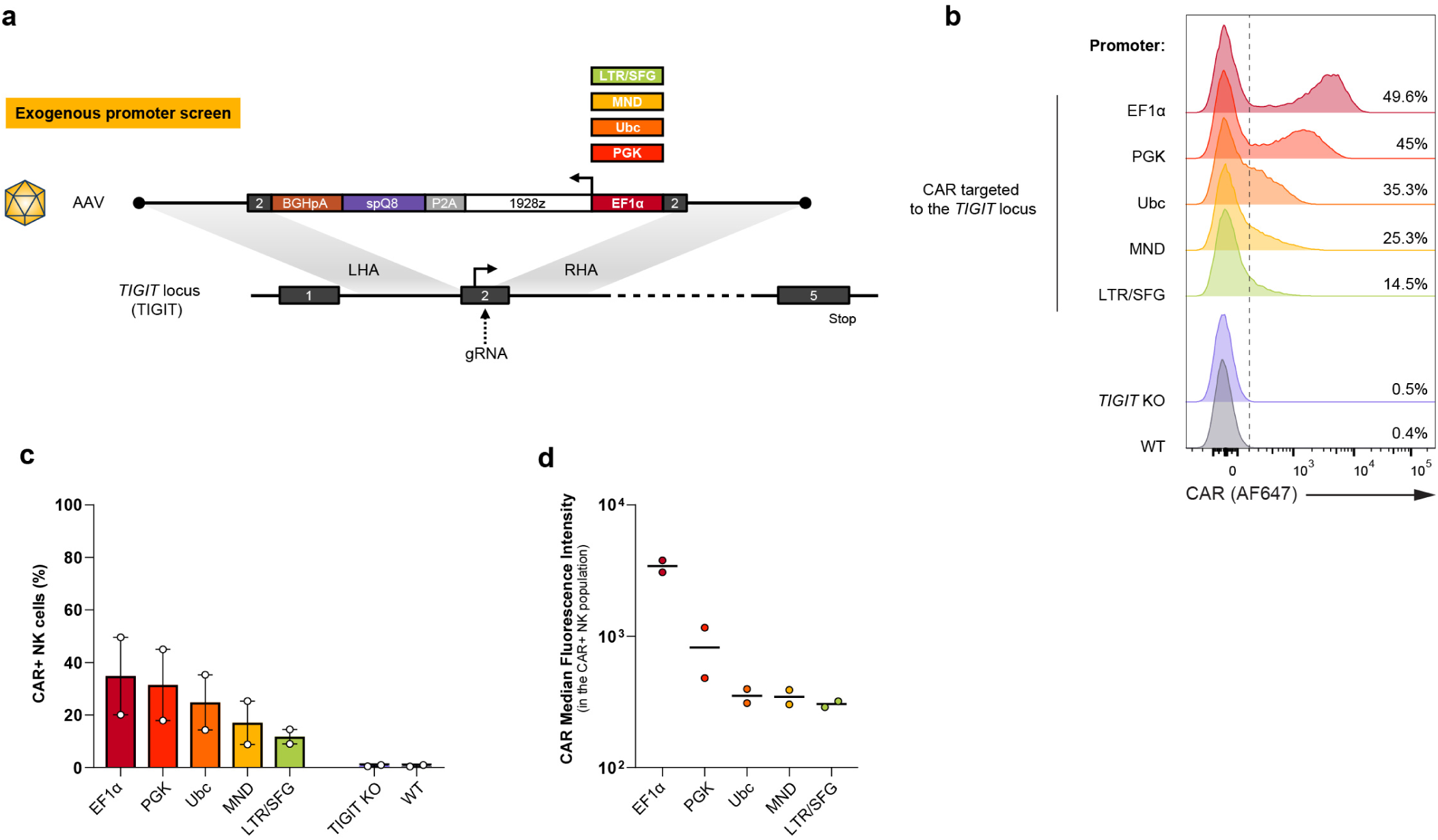
CAR expression level can be controlled using different exogenous promoters. **a**, Different exogenous promoters were compared for regulating a CAR targeted at the *TIGIT* locus (disruptive design) in primary NK cells: EF1α, PGK, Ubc, MND and LTR/SFG. **b**, Representative flow cytometry data acquired 5 d after editing. The percentage of CAR^+^ NK cells is indicated for each promoter. **c-d**, CAR^+^ percentage in NK cells (**c**) and CAR median fluorescence intensity in CAR^+^ NK cells (**d**) for *n*=2 different donors in 2 independent experiments.

**Extended Data Figure 8.**
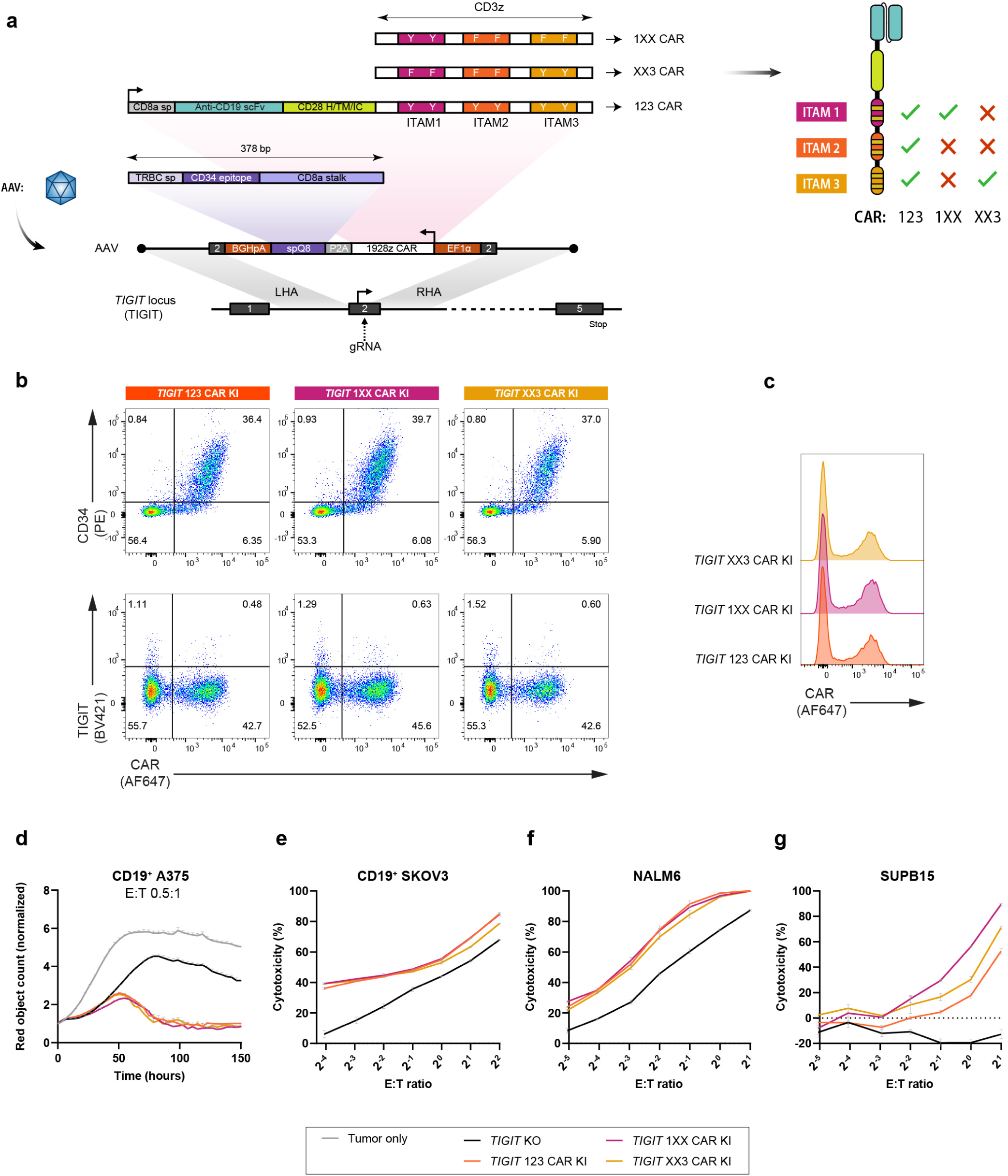
Targeted integration of CAR ITAM variants at the *TIGIT* locus. **a**, Three variants of the 1928z CAR, based on Feucht *et al.*, 2019^45^, were integrated at the *TIGIT* locus. The 123 CAR corresponds to the regular 1928z CAR, with three functional ITAM in the CD3z domain. 1×X and XX3 variants have only one functional ITAM at position 1 and 3, respectively, by mutation of the appropriate tyrosine residues. The CAR transgene also has a spQ8 marker (adapted from Philip *et al.*, 2014^46^) integrated in a bicistronic fashion. spQ8 is composed of a TRBC signal peptide, a CD34 epitope, and a CD8A stalk. **b**, Representative flow cytometry data showing the expression of the CAR and the spQ8 marker after targeted integration at the *TIGIT* locus. **c**, Histogram of CAR expression for the 3 CAR variants. **d-g**, *In vitro* cytotoxicity assays against different cell lines expressing the CD19 antigen. The Incucyte cytotoxicity assay against CD19^+^ A375 cell line was performed at a starting 0.5:1 E:T ratio (**d**) in presence of IL-2. Luciferase-based cytotoxicity assays were performed against CD19^+^ SKOV3 (**e**), NALM6 (**f**) and SUPB15 (**g**). The cytometry and functional data presented in this figure are representative of 2 independent donors.

**Extended Data Figure 9.**
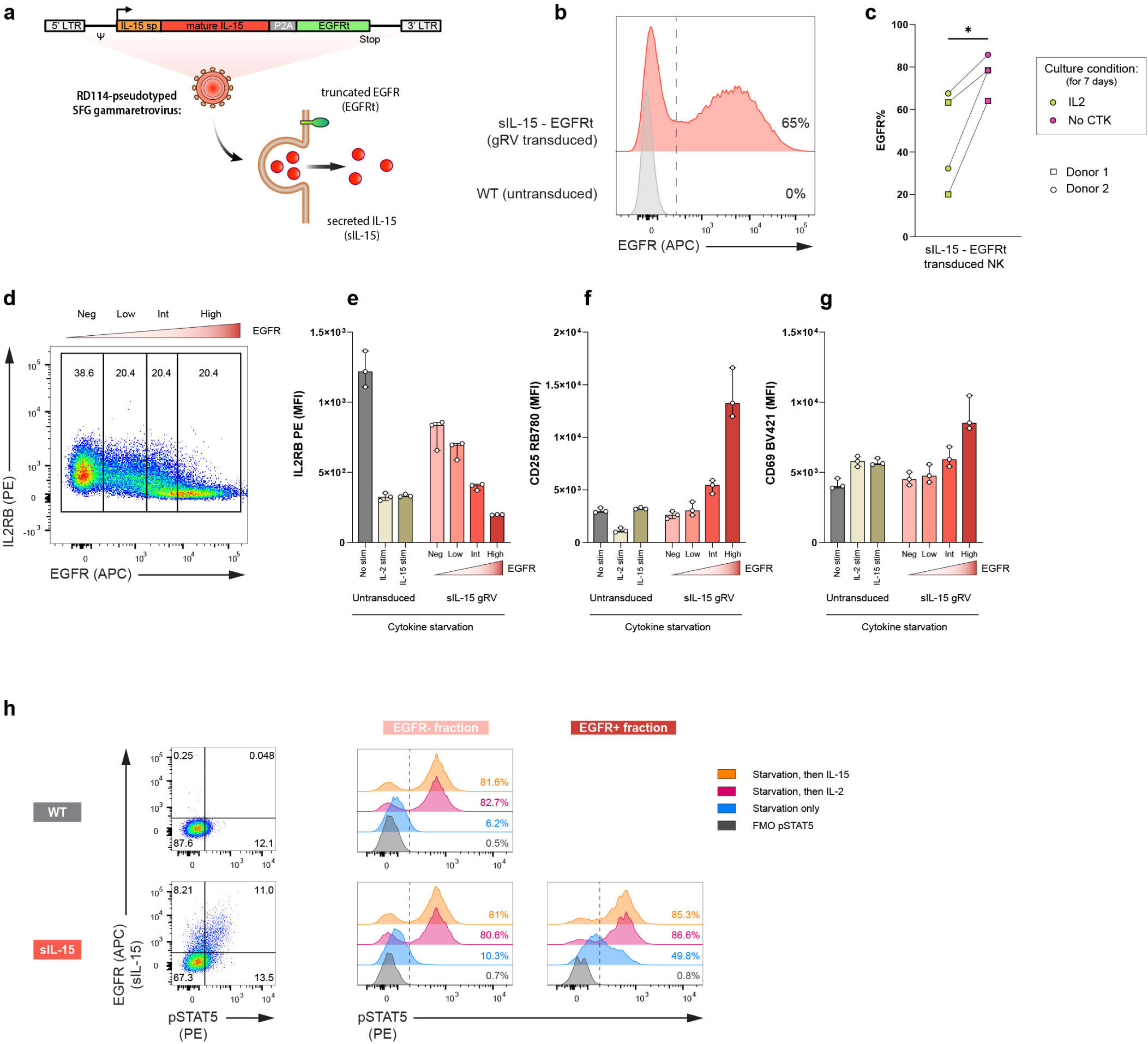
Constitutive expression of soluble IL-15 using a retrovirus vector in primary NK cells. **a**, Schematic of the RD114-pseudotyped gRV vector used for constitutive expression of soluble IL-15 (sIL-15) along with a truncated EGFR (EGFRt) membrane marker. **b**, Representative flow cytometry data showing transduction efficiency achieved with this vector. **c**, Enrichment of the EGFR percentage when sIL-15 expressing NK cells are cultured without cytokine compared to regular culture condition (IL-2 1000 U/mL) over a period of 7 d. n=2 independent donors transduced with a regular or low dose (1/10th of the regular dose) of gammaretrovirus. **d-g**, Untransduced and sIL-15 transduced NK cells cultured with regular IL-2 concentration were washed and starved from cytokines for 15 h and then subjected to flow cytometry for IL2RB (**d-e**), CD25 (**f**) and CD69 (**g**) mean fluorescence intensity (MFI) assessment. EGFR-positive cells were split in 3 tiers of equal size (Low, Int, High) (**d**). Controls included untransduced T cells stimulated with IL-2 or IL-15 for 2 h. *n*=3 technical replicates. Bars represent the median with error bars representing IQR. **h**, pSTAT5 phosphflow was performed in NK cells starved from cytokines for 15 h eventually followed by IL-2 (1000 U/mL) or IL-15 (10 ng/mL) stimulation for 30 min. pSTAT5 expression was assessed in the EGFR^-^ and EGFR^+^ fractions.

