## Supplementary information for "Site-specific genome engineering of primary human natural killer cells for programmable anti-tumor function"

### Supplementary Figures

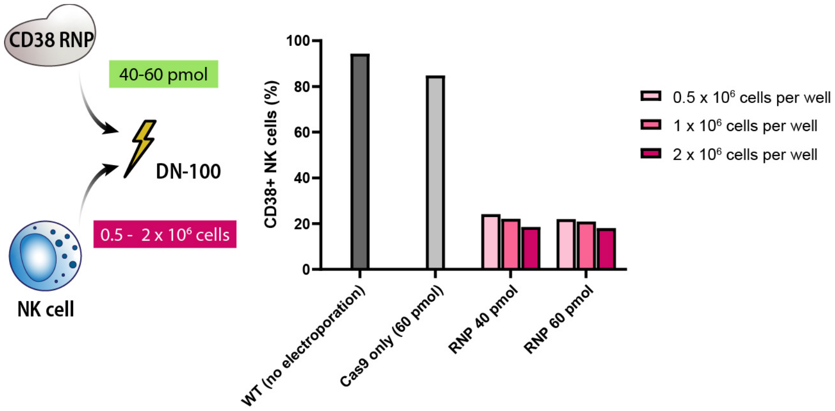


**Supplementary Figure 1**

**Optimization of electroporation parameters for *CD38* KO in primary NK cells - impact of cell number and RNP amount (related to Fig. 1)**

Primary NK cells were electroporated using the DN-100 pulse code with different amounts of cells (0.5 x $10^6$ to 2 x $10^6$) and *CD38* RNP (40 pmol to 60 pmol) per electroporation well. Flow cytometry was conducted 4 d after editing. Data represent the percentage of $CD38^+$ cells among NK cells.

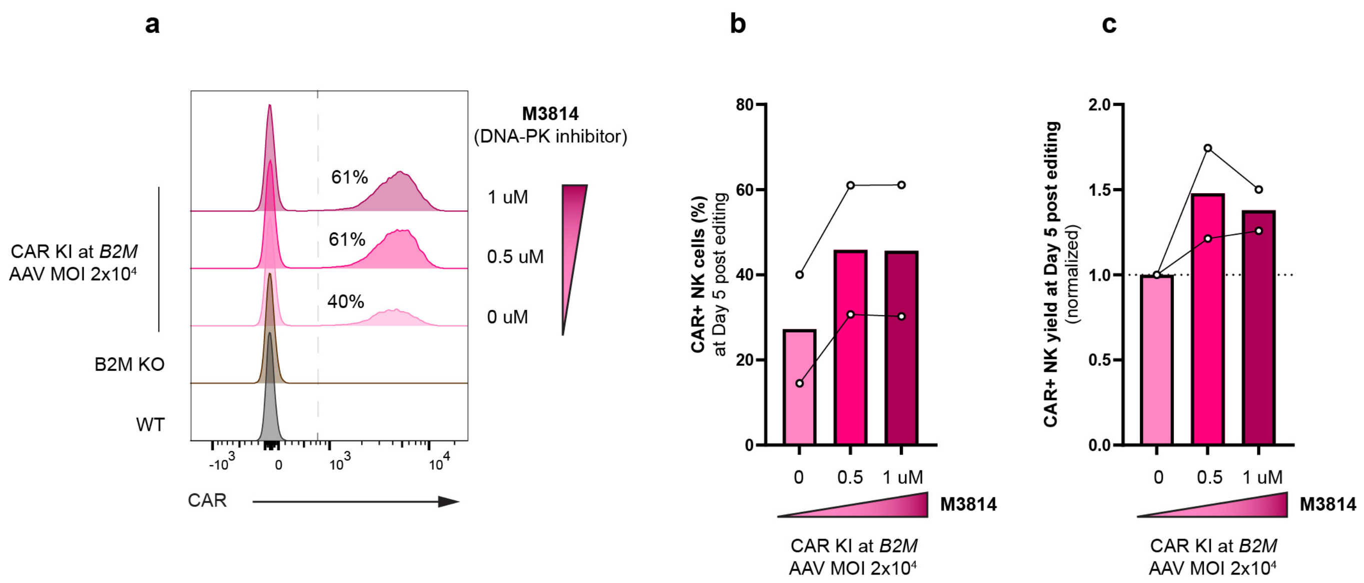


**Supplementary Figure 2**

**Increased knock-in efficiency by using the M3814 NHEJ inhibitor**

Targeted integration of a CAR at *B2M* (rescue design) was the editing model used to assess the impact of M3814, a DNA-PK inhibitor, on transgene targeted integration efficiency. **a**, Representative flow cytometry histogram showing the percentage of $CAR^+$ cells without or with two concentrations of M3814 during AAV transduction (0.5 and 1 μM). A low AAV MOI ($2 \times 10^4$) was intentionally used for these experiments. **b**, $CAR^+$ percentage in NK cells assessed by flow cytometry 5 d after editing. **c**, Overall editing yield ($CAR^+$ NK cells count), normalized to the condition without M3814, assessed 5 d after editing. Data are shown as mean. *n*=2 different donors (in 2 different experiments).

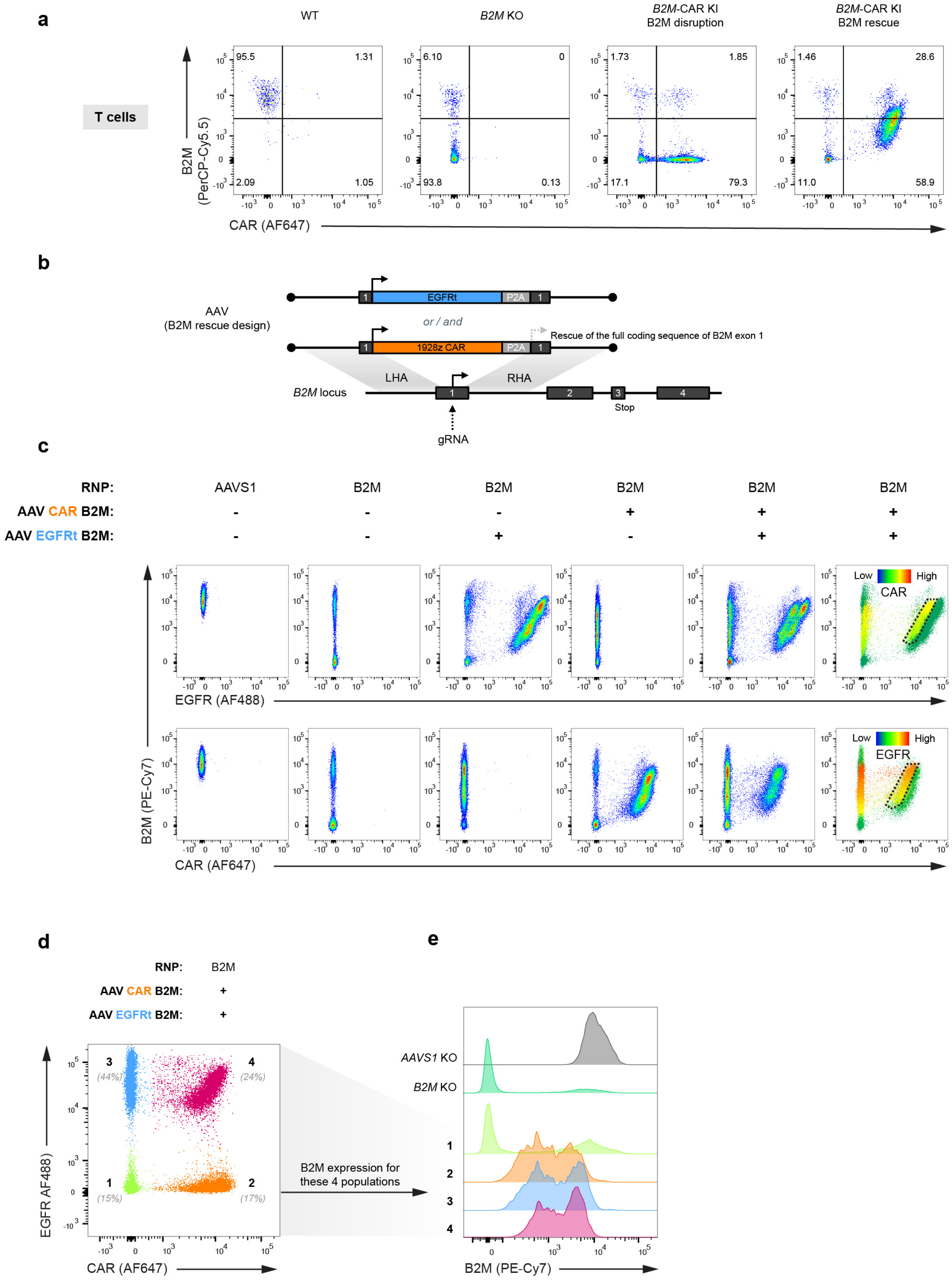
a
WT
B2M KO
B2M-CAR KI
B2M disruption
B2M-CAR KI
B2M rescue
T cells
B2M
(PerCP-Cy5.5)
95.5
1.31
2.09
1.05
6.10
0
93.8
0.13
1.73
1.85
17.1
79.3
1.46
28.6
11.0
58.9
CAR (AF647)
b
AAV
(B2M rescue design)
EGFRt
P2A
or / and
1928z CAR
Rescue of the full coding sequence of B2M exon 1
LHA
RHA
B2M locus
Stop
gRNA
c
RNP:
AAVS1
B2M
AAV CAR B2M:
AAV EGFRt B2M:
Low
High
CAR
EGFR
EGFR (AF488)
B2M (PE-Cy7)
CAR (AF647)
d
RNP:
B2M
AAV CAR B2M:
AAV EGFRt B2M:
(44%)
(24%)
(15%)
(17%)
EGFR AF488
CAR (AF647)
B2M expression for
these 4 populations
e
AAVS1 KO
B2M KO
B2M (PE-Cy7)

**Supplementary Figure 3**

**Intermediate B2M expression upon rescue is not explained by monoallelic transgene knock-in (related to Fig. 3)**

**a**, Representative flow cytometry data of primary T cells edited using the transgenes shown in **Fig. 3a**, demonstrating that intermediate B2M expression upon rescue is not NK cell specific. **b**, Schematic of the editing strategy for *B2M*-CAR KI or/and *B2M* EGFRt KI using a *B2M* rescue design. **c**, Flow cytometry data assessment at 5 d post-editing shows the respective expression of B2M, EGFR, and CAR for the different editing conditions shown in the upper rows. *AAVS1* KO cells were used as control. Flow panels on the right display a color scale for CAR and EGFR expression, to identify $EGFR^+$ and $CAR^+$ dual KI cells (area highlighted with a dotted black line). **d**, For the double AAV transduction (EGFRt and CAR simultaneous KI at *B2M* with a *B2M* rescue design), four subpopulations were identified based on their expression of EGFR and CAR. **e**, For the subpopulations defined in (**d**), B2M expression is represented as a flow cytometry histogram, with *AAVS1* KO and *B2M* KO cells used as positive and negative controls, respectively.

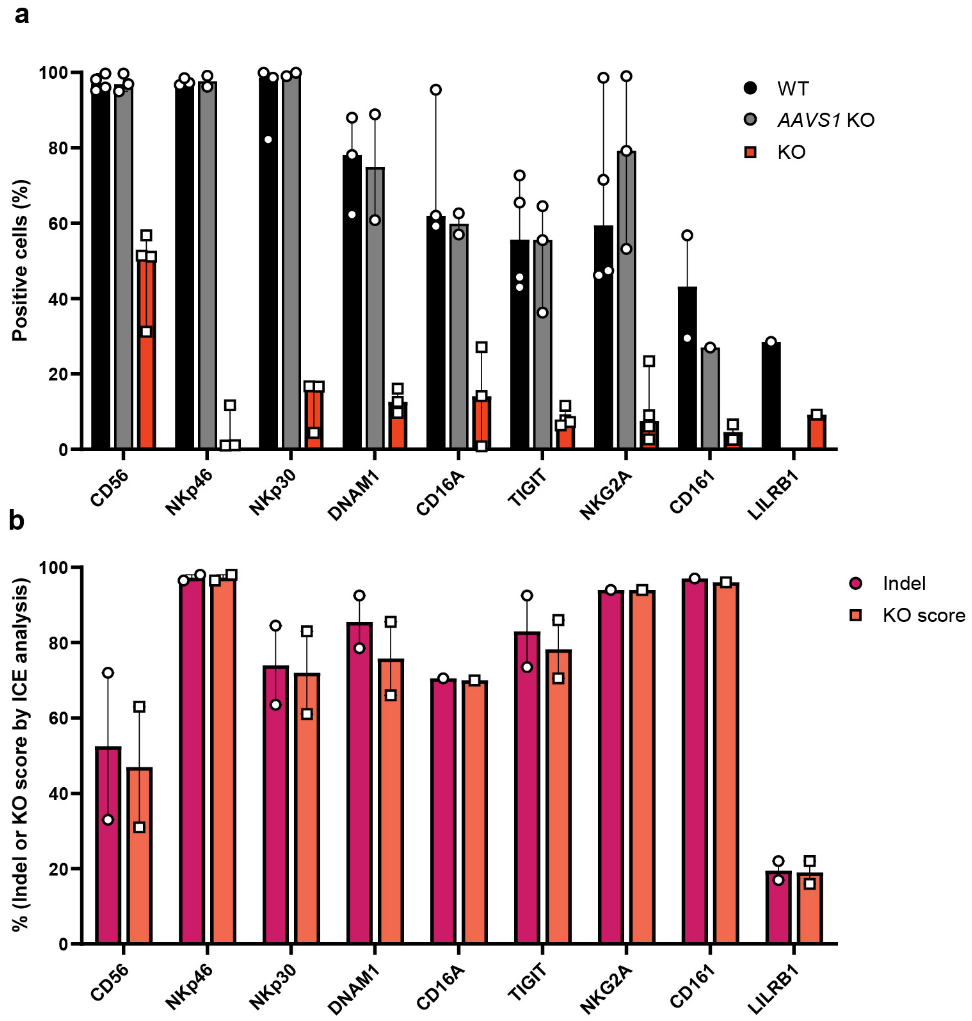


**Supplementary Figure 4**

**Knockout efficiency at different NK-relevant loci evaluated at the protein and the genomic levels (related to Fig. 4)**

**a**, Flow cytometry evaluation of KO efficiency at different loci. The graph represents the percentage of cells positive for the marker encoded by the target locus indicated on the x-axis. WT NK cells are used as a control for baseline expression of a given marker, as well as *AAVS1* KO NK cells when available. Data were compiled from *n*=4 different donors (4 independent experiments). **b**, Evaluation of genomic editing efficiency by ICE analysis after KO at different loci. Both indel percentage (purple) and KO score (orange) are shown. Each point represents the mean of the scores generated by ICE with both forward and reverse Sanger sequencing for a given donor. Data are pooled from 2 different donors. The data in (**a**,**b**) are presented as individual values, median and range.

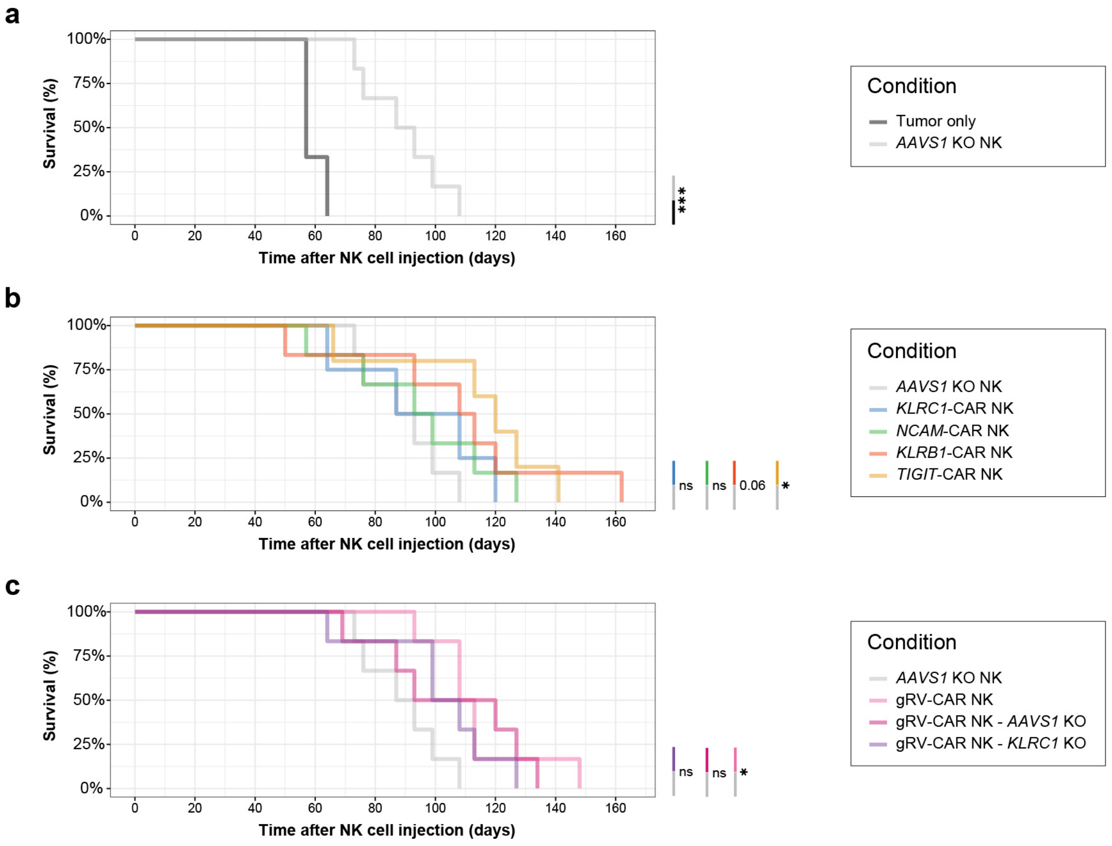


**Supplementary Figure 5**

**Split survival analysis in the SKOV3 xenograft mouse model for the screening of different loci for CAR targeted integration (related to Fig. 4)**

The curves in this figure are extracted from **Fig. 4i** and split in different panels for higher clarity and statistical analysis between treatment conditions. **a**, Comparison of *AAVS1* KO NK versus absence of treatment. **b**, Comparison of CAR NK cells engineered by targeting a CAR to different loci. **c**, Comparison of CAR NK conditions engineered by retrovirus transduction. *P*-values are from a log-rank test.

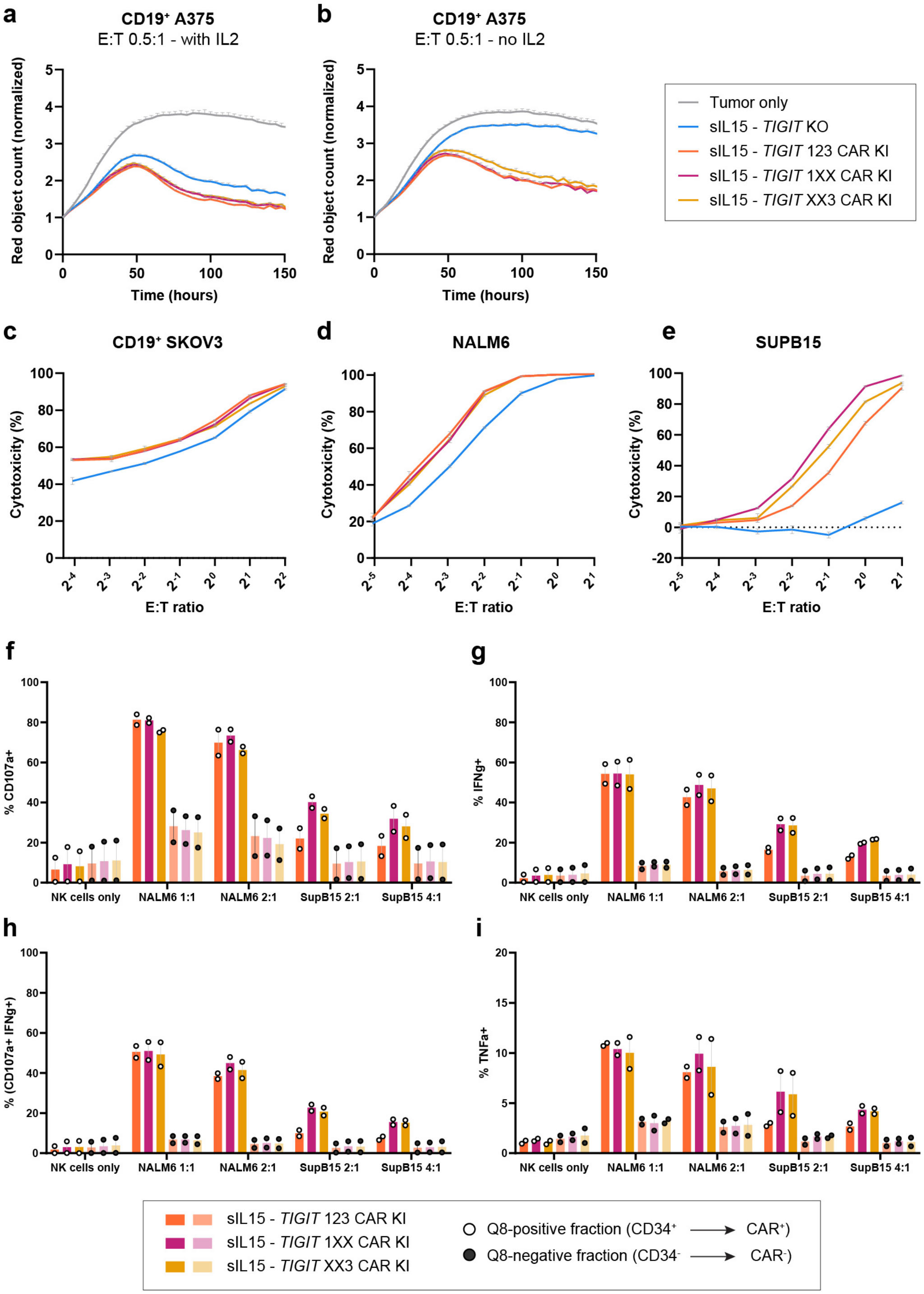


**Supplementary Figure 6**

***In vitro* functional characterization of CAR NK cells with targeted integration of CAR ITAM variants at the *TIGIT* locus combined with retroviral integration of a sIL-15 transgene.**

**a-e**, *In vitro* cytotoxicity assays against different cell lines expressing CD19. An Incucyte cytotoxicity assay against CD19+ A375 cell line was performed at a 0.5:1 E:T starting ratio with (**a**) or without (**b**) IL-2 in the coculture medium. Luciferase-based cytotoxicity assays were performed against $CD19^+$ SKOV3 (**c**), NALM6 (**d**) and SUPB15 (**e**). The cytotoxicity data in this figure are representative of $n$=2 (**c,e**) and $n$=3 (**a,b,d**) independent donors. **f-i**, Combined degranulation and cytokine production assay (interferon-γ and TNF-α) after 6 h of stimulation by NALM6 or SUPB15 target cells. For each cell line, two different E:T ratios were screened (1:1 and 2:1 for NALM6, 2:1 and 4:1 for SUPB15). The E:T ratio was based on the total number of NK cells for both cell lines. Percentage of $CD107a^+$ cells (**f**), $IFN\gamma^+$ cells (**g**), $CD107a^+$ $IFN\gamma^+$ double-expressor cells (**h**) and $TNF\text{-}\alpha^+$ cells (**i**) are represented for the three ITAM variants. The results are further divided between the $Q8^+$ and $Q8^-$ fractions using Q8 as a marker of CAR transgene integration.

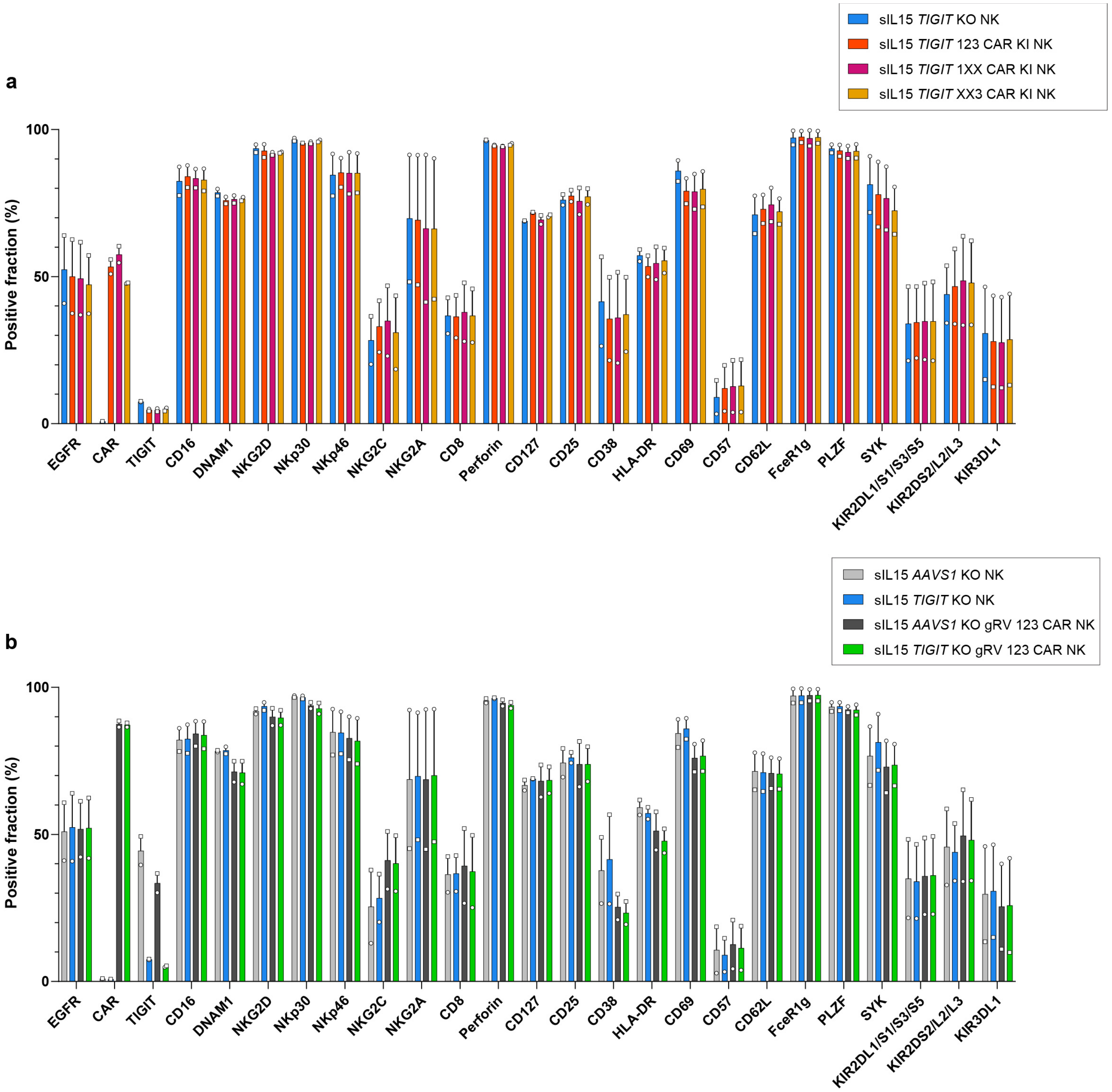


**Supplementary Figure 7**

**Flow cytometry phenotyping of different NK cell conditions (related to Fig. 5)**

High dimensional NK cell phenotyping by spectral flow cytometry was performed for different NK cell conditions in 2 independent donors (circle or square). **a**, Comparison of NK cells transduced with a retrovirus for sIL-15 constitutive expression (EGFRt marker) and targeted integration at *TIGIT* of an anti-CD19 CAR with three ITAM variants (123, 1XX, XX3). **b**, Comparison of NK cells transduced with a retrovirus for sIL-15 constitutive expression (EGFRt marker) and transduced or not with a second retrovirus for an anti-CD19 CAR expression, with either *AAVS1* or *TIGIT* KO. *n*=2 independent donors.

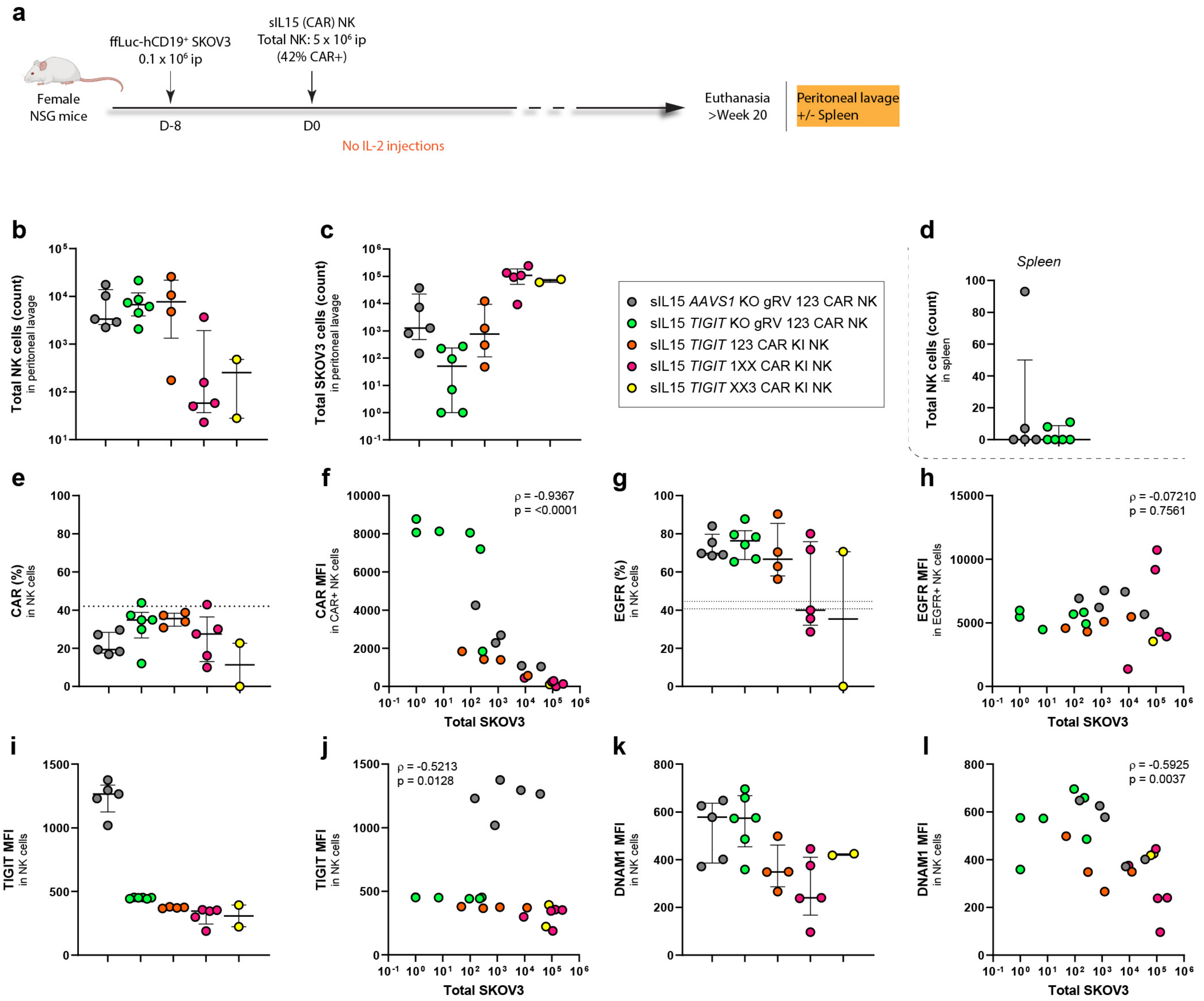


**Supplementary Figure 8**

**Flow cytometry quantification and phenotyping of NK cells recovered from peritoneal lavage in a mouse xenograft model of ovarian cancer (related to Fig. 5i-n)**

**a**, Schematic experimental design. Euthanasia and peritoneal lavage were performed in surviving animals after week 20 post NK injection. **b**, Total NK cell counts in the peritoneal lavage. **c**, Total SKOV3 cell counts in the peritoneal lavage. **d**, Total NK cell counts in the spleen for a subset of animals (CAR gammaretrovirus conditions). **e-f**, $CAR^+$ percentage in NK cells from the peritoneal lavage (**e**) and the corresponding CAR MFI according to the SKOV3 count (**f**). The dotted line corresponds to the $CAR^+$ percentage at the time of infusion in mice. **g-h**, $EGFR^+$ percentage in NK cells from the peritoneal lavage (**g**) and the corresponding EGFR MFI according to the SKOV3 count (**h**). The dotted lines correspond to the range of the $EGFR^+$ percentage at the time of infusion in mice. **i-j**, TIGIT MFI in NK cells from the peritoneal lavage for the different treatment groups (**i**) and according to the SKOV3 count (**j**). **k-l**, DNAM1 MFI in NK cells from the peritoneal lavage for the different treatment groups (**k**) and according to the SKOV3 count (**l**). Bars represent the median and error bars represent interquartile range. For (**f,h,j,l**), Spearman correlation *rho* was reported with the corresponding two-tailed *p*-value.

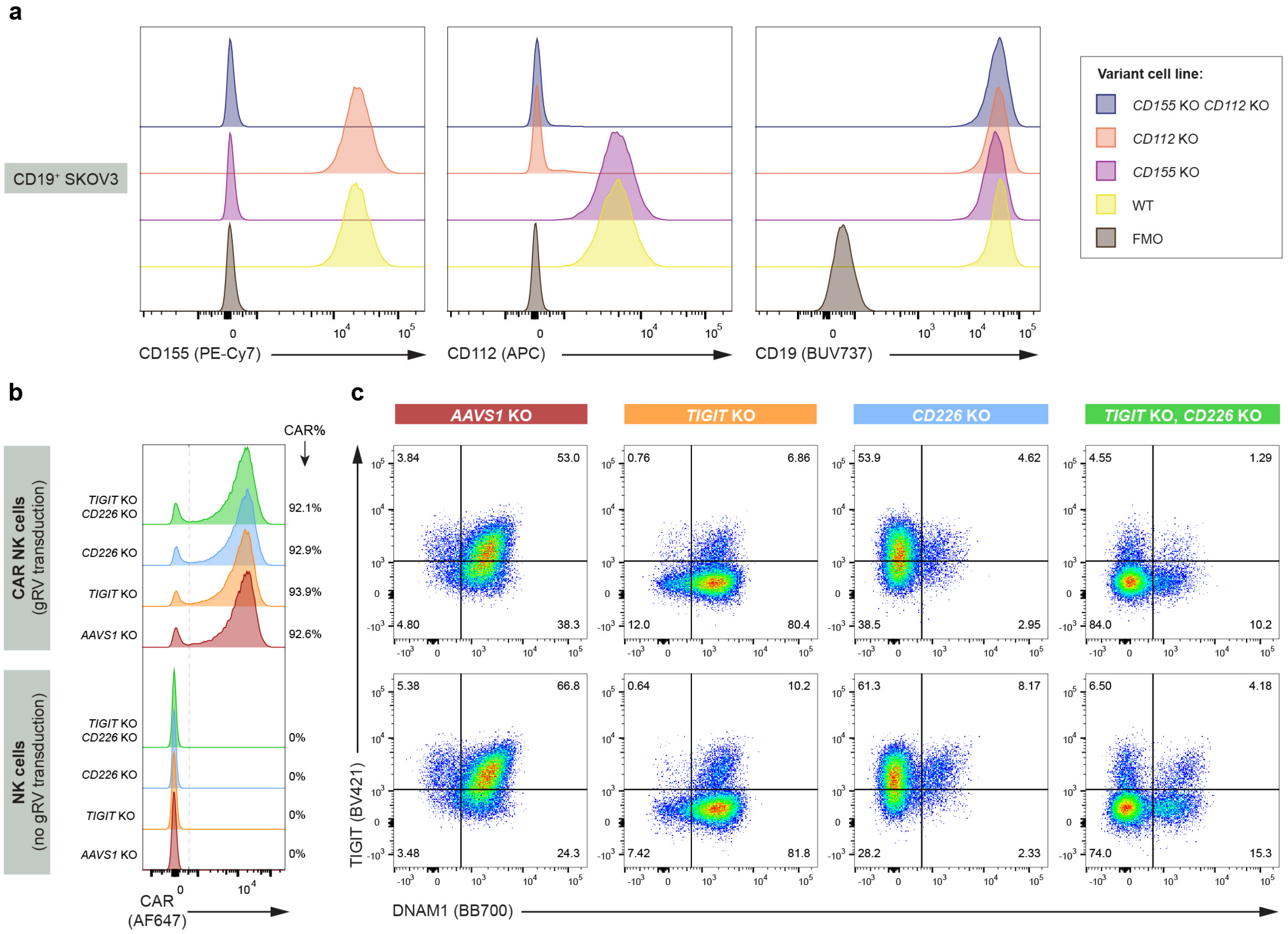


**Supplementary Figure 9**

**CD19$^{+}$ SKOV3 variants with targeted deletion of CD155 and/or CD112 - related to Supplementary Fig. 10.**

**a**, The CD19$^{+}$ SKOV3 cell line was engineered by CRISPR/Cas9 for specific deletion of *PVR* (CD155), *NECTIN2* (CD112), or both. Flow cytometry was performed on the resulting polyclonal cell lines to assess CD155, CD112, and CD19 surface expression. **b-c**, Primary NK cells were transduced (upper panels) or not (lower panels) with a gRV vector for anti-CD19 CAR transgene integration, followed by targeted deletion of *TIGIT* (TIGIT), *CD226* (DNAM1), or both (receptors for the CD155 and CD112 ligands). Deletion of *AAVS1* was used as control. Representative flow cytometry data showing the expression of the CAR (**b**), TIGIT (**c**) and DNAM1 (**c**) for the different NK conditions.

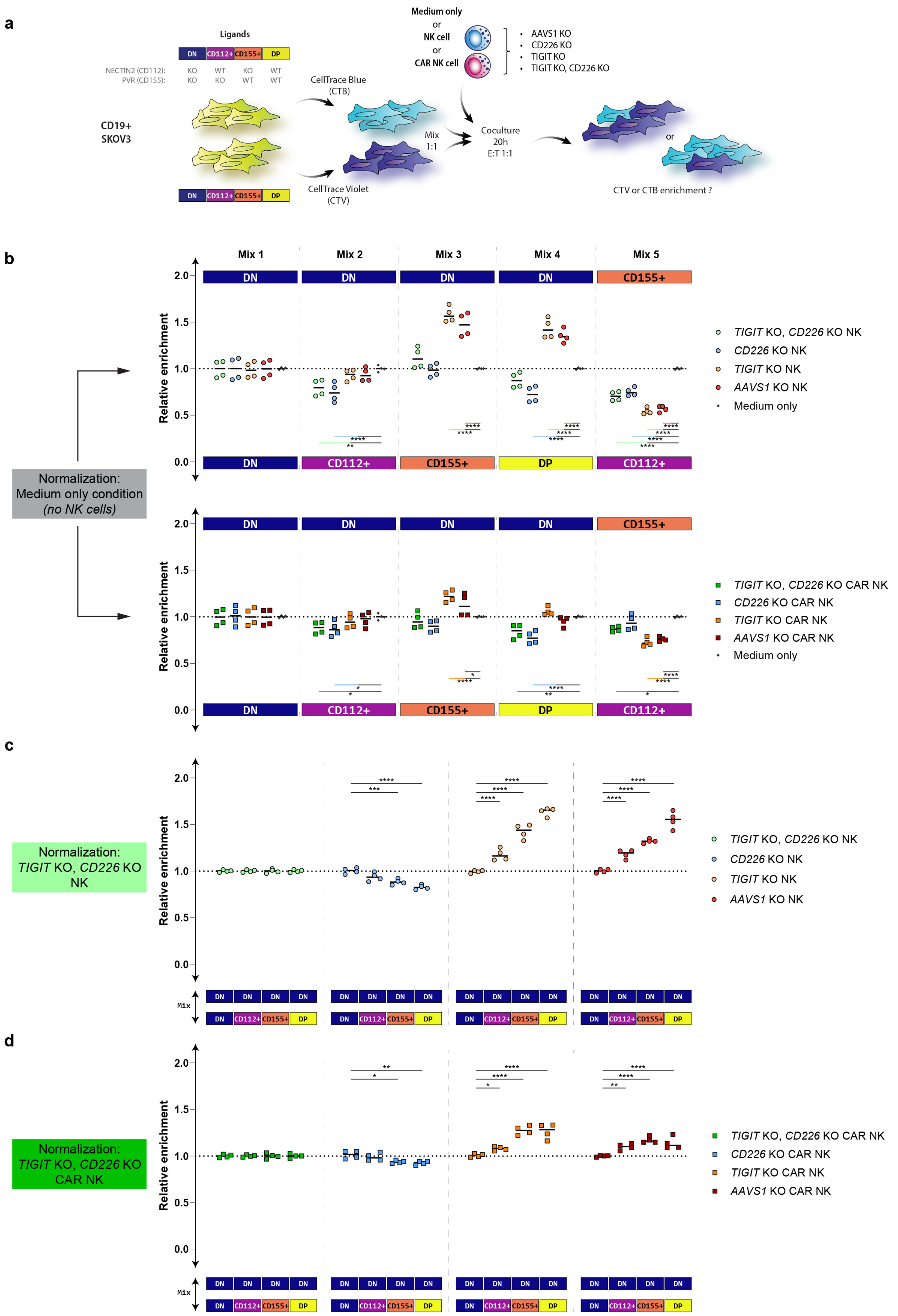
a
Ligands
DN CD112+ CD155+ DP
NECTIN2 (CD112): KO WT KO WT
PVR (CD155): KO KO WT WT
CD19+ SKOV3
CellTrace Blue (CTB)
CellTrace Violet (CTV)
Mix 1:1
Medium only or NK cell or CAR NK cell
AAVS1 KO
CD226 KO
TIGIT KO
TIGIT KO, CD226 KO
Coculture 20h E:T 1:1
or
CTV or CTB enrichment ?
b
Mix 1
Mix 2
Mix 3
Mix 4
Mix 5
Relative enrichment
Normalization: Medium only condition (no NK cells)
TIGIT KO, CD226 KO NK
CD226 KO NK
TIGIT KO NK
AAVS1 KO NK
Medium only
TIGIT KO, CD226 KO CAR NK
CD226 KO CAR NK
TIGIT KO CAR NK
AAVS1 KO CAR NK
Medium only
c
Normalization: TIGIT KO, CD226 KO NK
Mix
d
Normalization: TIGIT KO, CD226 KO CAR NK
Mix

**Supplementary Figure 10**

**Competitive cytotoxicity assay against CD19$^+$ SKOV3 cells expressing different combinations of CD155 and CD112 ligands.**

**a**, Schematic of the experimental design. Different mixes of target CD19$^+$ SKOV3 cells expressing or not CD155 and CD112 ligands (**Supplementary Fig. 9a**) were submitted to NK cell killing after labelling by either CellTrace Blue (CTB) or CellTrace Violet (CTV). Coincubation lasted 20 h with a starting effector to target ratio (E:T) of 1:1. (CAR) NK cells deleted for *CD226*, *TIGIT* or both were used as effectors, with *AAVS1* KO NK cells used as controls. At the end of the assay, the relative enrichment of CTB$^+$ versus CTV$^+$ surviving target cells was assessed by flow cytometry. **b**, Relative enrichment values were normalized by the corresponding "Medium only" condition for the same tumor mix. The upper panel corresponds to untransduced NK cells whereas the lower panel corresponds to CAR NK cells. **c-d**, Relative enrichment values were then normalized by the value of the corresponding *TIGIT* KO, *CD226* KO (CAR) NK cell condition for the same tumor mix. Untransduced (**c**) and CAR-transduced (**d**) NK cells are presented separately. $n$=4 technical replicates per condition. DN: Double negative; DP: double positive. $P$-values are from two-way ANOVA test with Dunnett's multiple comparison tests against the Medium only condition (**b**) or the DN/DN control mix (**c,d**). *, $p \leq 0.05$; **, $p \leq 0.01$; ***, $p \leq 0.001$; ****, $p \leq 0.0001$.

### Supplementary Tables

| **Locus** | **gRNA sequence** (5' to 3') | **F primer sequence** | **R primer sequence** |
|---|---|---|---|
| ***CD38*** **(gRNA1)** | GUUGGGCUCUCCUAGAGAGC | ATTGGTGTAACCAGCCACGG | CACGGAAGATGCCCGGTT |
| ***CD38*** **(gRNA2)** | CUCCUAGAGAGCCGGCAGCA | ATTGGTGTAACCAGCCACGG | CACGGAAGATGCCCGGTT |
| ***CD7*** **(ABE)** | CCCUACCUGUCACCAGGACC | ACCGTGTCTTTGGCGACAT | ACATGTAGGAGGGAGGGTCC |
| ***AAVS1*** **(ABE)** | GUCACCAAUCCUGUCCCUAG | CTCTGGCTCCATCGTAAGCA | CGTTCTCCTGTGGATTCGGG |
| ***B2M*** | GGCCACGGAGCGAGACAUCU | ACAGCAAACTCACCCAGTCT | AAACTTTGTCCCGACCCTCC |
| ***TIGIT*** | UCCUCCUGAUCUGGGCCCAG | GGCACTACACAGATGCCCTT | TCCTGCTCCTTCTTGTGCTG |
| ***NCAM*** | GGAUAUUGUUCCCAGCCAGG | CCACACAACCTCCTCCCTTC | GAGCCAGTGCTGTTCTCCAT |
| ***NCR1*** | GAACUCACCGACGCAGAGCA | CACAGTCAGCTCTGGGTACG | CCTGGCAACAGATGGTCACT |
| ***NCR3*** | CAAGAUGAGCAACAGCAUCC | CTCCTGGAGGCTTGTTCTGG | CCTCACAGTGGGTGACACAG |
| ***CD226*** | GCUCUUCUUCAUGUAUACAG | ATAAGCCGACTGGTTCCCCG | AGGGCTTCCTTATGACCATGC |
| ***KLRC1*** | ACUGCAGAGAUGGAUAACCA | TTGCTTCTCATTGCCCCAGC | CCCAAGCTGCACATCCTAGA |
| ***KLRB1*** | AAUUAAAGCCACUUACCCCG | TGTGGGTGGGGACAAAAGAG | GCCCAGGGTAGCCTAAAGTG |
| ***FCGR3A*** | UCGAGCACCCUGUACCAUUG | TGCTGATGTGGGGTTAGCAG | TGGAGAGGTTTGTCTGGCAC |
| ***LILRB1*** | CUGGCCCCCCUGACACCUGA | TGAGTCTGTCCCCAGCTCTT | AAATGTGGCTCAGACCACCC |
| ***AAVS1*** | GGGCCACUAGGGACAGGAU | N/A | N/A |
| ***PVR*** | GAUGUUCGGGUUGCGCGUAG | N/A | N/A |
| ***NECTIN2*** | CAAGUGCUACCCGAGGUGCG | N/A | N/A |

**Supplementary Table 1**

**List of sgRNA used and corresponding primers for ICE analysis**

| Antigen | Clone | Color | Supplier |
|---|---|---|---|
| B2M | 2M2 | PE-Cy7, PerCP-Cy5.5 | BioLegend |
| B7-H6 | 875001 | PE | R&D Systems |
| Calreticulin | 681233 | AF647 | R&D Systems |
| CD3 | UCHT1 | BUV395, BV605 | BD Biosciences |
| CD3 | SK7 | Spark NIR 685 | BioLegend |
| CD8 | SK1 | Spark Blue 550 | BioLegend |
| CD16 | 3G8 | AF488, BUV496 | BioLegend |
| CD19 | HIB19 | BV510 | BioLegend |
| CD19 | SJ25C1 | BUV737 | BD Biosciences |
| CD25 | BC96 | RB780, BV650 | BD Biosciences |
| CD34 | QBEND/10.rMab | PE, BV421 | BD Biosciences |
| CD38 | HIT2 | BV510 | BioLegend |
| CD45 | 2D1 | APC-Cy7, PerCP | BD Biosciences |
| CD48 | BJ40 | PE-Cy7 | BioLegend |
| CD54 | HA58 | BV421 | BD Biosciences |
| CD56 | HCD56 | BV711 | BioLegend |
| CD56 | NCAM16.2 | BUV737 | BD Biosciences |
| CD57 | QA17A04 | PE | BioLegend |
| CD57 | NK-1 | BUV395 | BD Biosciences |
| CD58 | TS2/9 | PE-Cy7 | BioLegend |
| CD62L | SK11 | BB700 | BD Biosciences |
| CD69 | FN50 | Pacific Blue, BV421 | BioLegend |
| CD107a | H4A3 | FITC | BD Biosciences |
| CD112 | TX31 | APC | BioLegend |
| CD127 | A019D5 | PE-Cy5 | BioLegend |
| CD155 | SKII.4 | PE-Cy7 | BioLegend |
| CD158a | DX9 | BV711 | BD Biosciences |
| CD161 | HP-3G10 | BV480, BV785, PE | BioLegend |
| DNAM1 | DX11 | BB700, BUV805 | BD Biosciences |
| EGFR | AY13 | AF488, APC, BV785 | BioLegend |
| Fcer1g | Polyclonal | FITC | Millipore |
| Mouse IgG, F(ab')$_2$ fragment specific | Polyclonal; F(ab')$_2$ Fragment (Goat Anti-Mouse) | AF647 | Jackson ImmunoResearch, Laboratories |
| G4S Linker | G4S (E7O2V) | PE, AF647 | Cell Signaling Technology, Inc. |
| HLA Class I | W6/32 | BV510 | BioLegend |
| HLA-E | 3D12 | BV421 | BioLegend |
| HLA-DR | G46-6 | BUV661 | BD Biosciences |
| IFNg | B27 | BV421 | BioLegend |
| IL2RB | TU27 | PE | BioLegend |
| KIR2DS2/L2/L3 | DX27 | BUV563 | BD Biosciences |
| KIR2DL1/S1/S3/S5 | HP-MA4 | APC/Fire 780 | BioLegend |
| KIR3DL1 | DX9 | BV711 | BD Biosciences |
| LILRB1 | GHI/75 | BUV615, PE | BD Biosciences |

| | | | |
|---|---|---|---|
| LLT-1 | 402659 | PE | R&D Systems |
| MICA/B | 6D4 | BV421 | BD Biosciences |
| NGFR | C40-1457 | BV421 | BD Biosciences |
| NKG2A | S19004C | AF647 | BioLegend |
| NKG2A | 131411 | BV650 | BD Biosciences |
| NKG2C | 134591 | BV480, PE | BD Biosciences |
| NKG2D | 1D11 | BV750 | BD Biosciences |
| NKp30 | P30-15 | PE, BV605 | BioLegend |
| NKp46 | 9E2 | PE-Cy7 | BioLegend |
| NKp46 | 9E2 | SB436 | Invitrogen |
| Perforin | dG9 | PerCP/Cy5.5 | BioLegend |
| PLZF | R17-809 | PE/CF594 | BD Biosciences |
| pSTAT5 | 47/Stat5 (pY694) | PE | BD Biosciences |
| SYK | 4D10.2 | APC | BioLegend |
| TIGIT | A15153G | BV421 | BioLegend |
| TNFa | MAb11 | PE-Cy7 | BioLegend |
| ULBP-1 | 170818 | PE | R&D Systems |
| ULBP-2/5/6 | 165903 | APC | R&D Systems |

**Supplementary Table 2**

**List of antibodies used for flow cytometry analysis.**
